# Stochastic fluctuations drive non-genetic evolution of proliferation in clonal cancer cell populations

**DOI:** 10.1101/2021.06.08.447479

**Authors:** Carmen Ortega-Sabater, Gabriel F. Calvo, Jelena Dinić, Ana Podolski-Renic, Milica Pesic, Víctor M. Pérez-García

**Affiliations:** Mathematical Oncology Laboratory (MOLAB), Universidad de Castilla-La Mancha, Ciudad Real, 13071, Spain; Institute for Biological Research Sinia Stanković, University of Belgrade, Despota Stefana 142, 11060, Belgrade, Serbia

**Keywords:** evolutionary dynamics, cancer, phenotype, stochastic, noise expression

## Abstract

Evolutionary dynamics allows us to understand many changes happening in a broad variety of biological systems, ranging from individuals to complete ecosystems. It is also behind a number of remarkable organizational changes that happen during the natural history of cancers. These reflect tumour heterogeneity, which is present at all cellular levels, including the genome, proteome and phenome, shaping its development and interrelation with its environment. An intriguing observation in different cohorts of oncological patients is that tumours exhibit an increased proliferation as the disease progresses, while the timescales involved are apparently too short for the fixation of sufficient driver mutations to promote an explosive growth. Here we discuss how phenotypic plasticity, emerging from a single genotype, may play a key role and provide a ground for a continuous acceleration of the proliferation rate of clonal populations with time. We address this question by combining the analysis of real time growth of non-small-cell lung carcinoma cells (line NCI-H460) together with stochastic and deterministic mathematical models that capture proliferation trait heterogeneity in clonal populations to elucidate the contribution of phenotypic transitions on tumour growth dynamics.

## 1. Introduction

Evolution is one of the central unifying concepts of biology and a driving force behind life, being a cornerstone of complex systems organization [1]. It is ubiquitous through the natural world from molecules to cells, organisms and populations and in fields as diverse as zoology, botany, microbiology and oncology. Evolutionary changes in the context of asexual reproduction are mainly driven by heritable somatic mutations and epigenetic changes, genetic drift and natural selection. Evolution theory has been classically grounded in genetics and Darwinian selection processes. In the light of evolution, tumour progression has often been explained by looking at the somatic changes of cancer cells [2,3]. However, from this viewpoint, we might be missing the bigger picture and incur in some assumptions that are sometimes in conflict with what is observed during the real course of the disease, including treatment failure and relapse [4].

There is a growing interest in studying the evolutionary forces behind cancer, and a number of important questions remain open. Increased attention has recently been given to intratumour heterogeneity as its potential role in the emergence of drug resistance and therapeutic outcome [5**?** –7]. Intratumour heterogeneity is the result of the clonal diversity within tumors [8]. It occurs at various levels, including the genome, transcriptome, proteome and phenome [9]. Research has mainly focused on mapping cancer genome instability and driver event mutations conferring a selective advantage to the affected cell clone. However, mutation-independent instability and noisy variability affecting transcription and translation of DNA have previously been reported in both prokaryotes and eukaryotes, even in homogeneous environments [10,13,14,39].

Stochasticity has been proposed as a generator of phenotypic heterogeneity in microorganisms, being especially useful under changing environmental conditions [15]. Similarly, initial states of cancer development involve colonization of novel environments and subsequent stressful conditions [16], which may actually modify traits heritability (understanding heritability as the relation between genetic variance for the trait and the phenotypic variance for the same trait). Thus, phenotypic variability is the result of a compendium of phenomena: the variations related to differences among genotypes, those associated with environmental changes and the ‘interaction variance’ which represents that some genotypes might respond to the environment in a different way than others [17]. Distinct phenotypic states frequently involve differences in functional cell properties and the proportion of these phenotypes has been related to cancer grade [22,23]. This resembles what occurs in other biological contexts, where a broader population composition, which comprises a higher phenotypic diversity, increases the odds for an adaptive response to external perturbations [24].

Understanding the growth patterns of tumours from a mathematical point of view has been the subject of many previous works (see for instance the following reviews and references therein [18– 21]). A fundamental question is whether by incorporating into the traditional conceptual representation of a tumor, viewed as an exponential-like increasing mass of proliferating cells subjected to deregulated molecular signalling pathways, the genotypic/phenotypic diversity may dramatically alter the quantitative models used to describe tumour growth. In this regard, scaling laws constitute an insightful way to characterize such growth patterns. Scaling laws are quantitative relationships of the form *Z* = *αV* ^*β*^ connecting an observable quantity of a complex system, say *Z*, with another measurable variable, say *V*, which in living systems is typically the volume or mass. In those laws *α* is a rate constant and *β* the scaling exponent.

Recent observations in both *in vivo* murine models and cohorts of cancer patients of different histologies have found a superlinear scaling law relating proliferation and tumour size [25]. Also, a longitudinal dynamics was observed implying a continuous acceleration of proliferation rates during tumour natural development, which is a dynamical counterpart of the scaling law. This fact was attributed initially to the tumour’s genetic evolutionary dynamics and supported with different mathematical modelling frameworks. However, a closer look at this interpretation raises a number of questions. The results presented in [25] included two studies in animal models that displayed accelerated tumour growth dynamics in the course of one month. Longitudinal volumetric data obtained from images of cancer patients with untreated brain metastases along the course of few months also showed similar growth patterns. Genetic changes seem to be necessary [26] but would not suffice to generate such accelerated tumour growth since they would require either an extremely high mutation rate, which would be restricted by cell viability, or a long time scale [27] that was not the case in the former results. For these reasons, neither the human data spanning typically few months, nor the animal models data, could be exclusively associated with tumour genetic changes because of the short time scales involved. This raises the question that we wish to explore in this paper: Could there be additional non-genetic evolutionary forces playing a role in accelerating tumour growth as observed in Ref. [25]?

The recognition of the role of phenotypic aspects in cancer evolutionary dynamics has elicited a progressive change in perspective from seeing cancer as a ‘genetic disease’ to a broader, ‘developmental’ perspective. Some analogies with embryonic development have been made and phenotypic plasticity in cancer has been biologically addressed in the literature of cancer stem cells [28]. Also, diapause-like states have been described as a survival mechanism against chemotherapy [29,30]. In this context, an initial clonal population will be subject to atypical evolutionary pressures in a stochastic or environmental-induced way that finally shape the population structure.

Fluctuations in proliferation rates of clonal cancer cell populations have been observed in cultures [31–33]. The repetition of the same cell-culture experiment leads to large variations in the outcome that cannot be attributted only to differences in the number of cells seeded initially. Some authors have previously accounted for these baseline variations in cell cycle duration among heterogeneous cancer cell populations [34,35] that could be indeed tightly linked to tumour response to therapy [36]. Phenotypic plasticity could affect any cell trait [37], but here we will focus on proliferation as a key phenotypic characteristic in cancers. Specifically, we study both *in vitro* and *in silico* the possibility that stochastic changes in the growth rate of clonal populations could lead to an evolutionary dynamic of that trait. Our main hypothesis is that small phenotypic changes resulting in either faster or slower proliferation could emerge as a result of noise-induced nongenetic variability. This may give rise to a variability in clonal populations that can provide the appropriate ground for selection and evolutionary dynamics. Figure 1 illustrates a particular setting of the general problem that we address here. Growth curves extracted from real time assessment of non-small-cell lung carcinoma (NCI-H460) cultures under different initial cell numbers (Fig. 1**a**) may reflect two distinct scenarios in which the cellular division time randomly varies (Fig. 1**b** and **c**). Associated to each of these scenarios there is a proliferation rate that may either display oscillations around some basal value or an increasing trend (Fig. 1**d** and **e**). By looking at the temporal window during which the cell culture is far from confluence we may determine, via discrete and continuous mathematical models, the underlying cell dynamics with the final goal of elucidating what is the contribution of phenotypic changes to the main driving force behind the accelerated tumour growth observed in human cancers [25].

**Figure 1.**
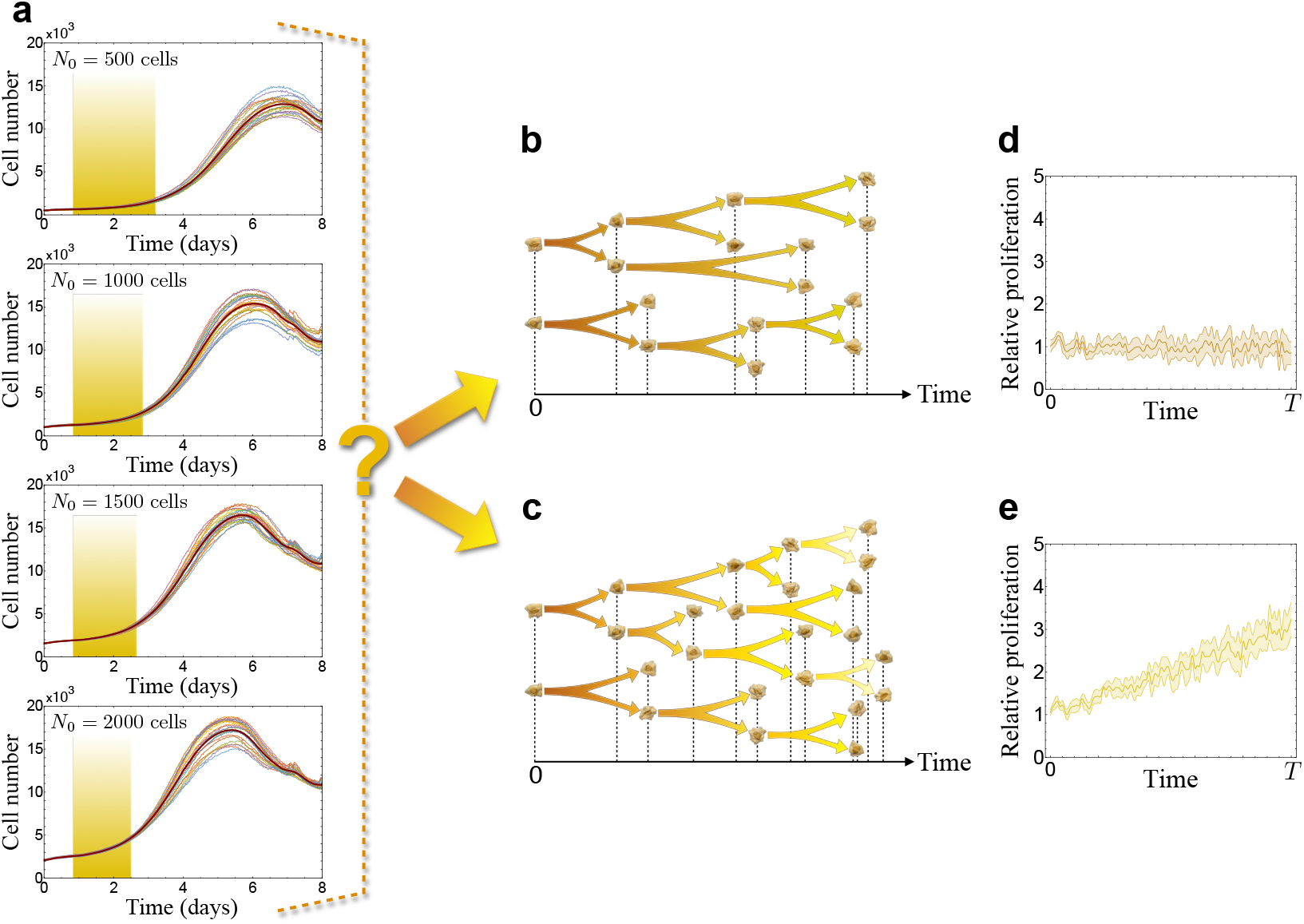
Stochastic fluctuations of cancer cell proliferation in two distinct scenarios. **a** Real time assessment of non-small-cell lung carcinoma cells (NCI-H460) with four different initial cell numbers seeded as explained in ‘Methods’ (*N*_0_ = 500, 1000, 1500, 2000 cells). The shaded windows indicate the selected time intervals where culture conditions were far from confluence. **b**-**e** Schematic representation of two possible scenarios were tumour cells divide randomly following either: **b** random choices of proliferation rates around a basal value or **c** An increasing mean proliferation rate. **d** and **e** The time-varying proliferation rate in both scenarios in units of the initial proliferation rate during a temporal window [0, *T*].

## 2. Results

To quantify the impact of phenotypic changes in proliferation on the growth dynamics of a clonal tumour cell population, we resorted to two mathematical models. The first one was based on a discrete simulator incorporating stochastic jumps between different proliferative states. The second one, consisting of a continuous reaction-diffusion parabolic equation, recapitulated the key aspects of the discrete model and allowed us to find explicit analytical formulas for the temporal dynamics of the total tumour cell number, together with the mean and the standard deviation in proliferation, that were subsequently fitted to the experimental data from the NCI-H460 cell line. We first considered *inheritable* phenotypic changes, meaning that when a cell is committed to mitosis, its progeny will be placed in the same proliferative state. In an alternative scenario, to explore the possibility of a *partial loss of inheritance* (and thus of partial reversibility), we also regarded the effect of adding a decay probability to the initial characteristic proliferation rate after the completion of a certain number of cell divisions.

### Stochastic fluctuations drive proliferation rate growth in clonal cancer cell populations with inheritable proliferative traits in-silico

First, we simulated the fully inheritable case in a time frame of *T* = 30 days by means of the discrete stochastic model presented in the Methods. This temporal frame was sufficiently short to discard relevant mutational events. An example of a typical outcome is shown in Fig. 2**a**. Even when the phenotypic transitions were fully symmetric, the system spontaneously drifted towards higher proliferation values, with the mean proliferation rate ⟨*ρ*⟩(*t*) reaching levels more than two times larger than the initial one *ρ*_*∗*_ [see inset Fig. 2**a**]. Also, a broadening in the phenotype landscape with time was apparent in the simulation results. In addition, a reaction-advection-diffusion model derived from the discrete stochastic model (see Methods) allowed us to reproduce these features in the same time frame, as depicted in Fig. 2**b**, together with the standard deviation rate [inset Fig. 2**b**]. Moreover, both models predicted that the total cell population, when plotted in a logarithmic scale, increased much faster than a simple exponential growth, as shown in Fig. 2**c**.

**Figure 2.**
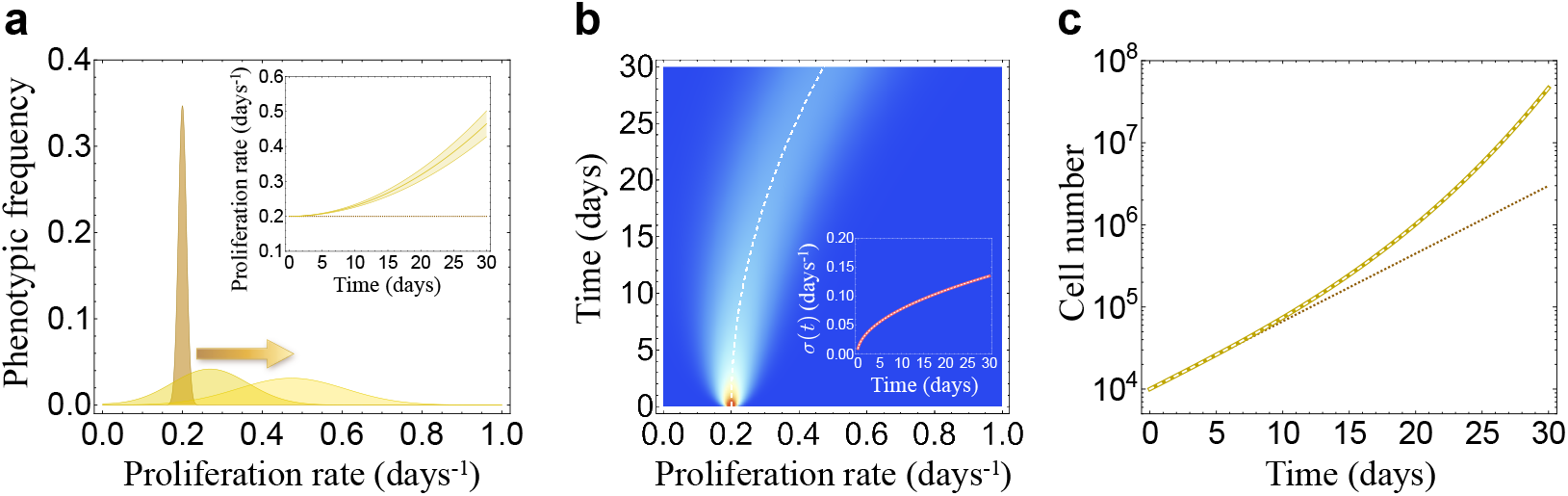
*In silico* evolutionary dynamics in the landscape of full phenotype inheritance. **a** Phenotypic frequency at times *t* = 0, 15, and 30 days (from left to right) of a typical run using the discrete stochastic model. The inset shows a time-dependence of the mean proliferation rate as an increasing broadening band, whereas the dotted line corresponds to a constant proliferation rate *ρ*_*∗*_ = 0.2 days^−1^. **b** Pseudocolor plot of the population cell density for *ρ* ∈ [0, 1] days^−1^ and *t* ∈ [0, 30] days computed from the continuous reaction-advection-diffusion model (4.3). The dashed white line indicates the mean proliferation rate ⟨*ρ*⟩(*t*) of the distribution as predicted by Eq. (2.5). The inset shows the standard deviation ⟨*σ*⟩(*t*) from the solution of (4.3) (thick reddish curve) and the analytical formula (2.6) (overlapping dashed white curve). **c** Dynamics of the total cell population *N* (*t*) from the solution of (4.3) (thick golden curve) and the analytical formula (2.4) (overlapping dashed white curve). The dotted line corresponds to the case of a purely exponential growth with a constant proliferation rate *ρ*_*∗*_. Numerical values used: For the discrete stochastic model *Γ* = 6.0 days^−1^, *M* = 101 nodes, *Δt* = 1 h, whereas the number of simulation runs was equal to 50. For the continuous reaction-advection-diffusion model, *D* = 3.0 × 10^−4^ days^−3^ and *v* = 0 days^−2^. For both the discrete and the continuous models, we used an initial Gaussian distribution centred around *ρ*_*∗*_ = 0.2 days^−1^ and initial standard deviation *σ*_*∗*_ = 0.01 days^−1^. Also, *µ* = 0.01 days^−1^, *N*_0_ = 10^4^ tumour cells, *ρ*_min_ = 0 day^−1^ and *ρ*_max_ = 1 day^−1^.

To determine the underlying explicit time dependences of all these distinctive characteristics, and hence gain further insight, we employed a reaction-advection-diffusion model [see Eq. (4.3) in Methods] in order to derive a complete set of ordinary differential equations for time-evolving average quantities. They are

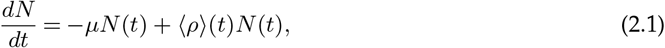

for the total cell number *N* (*t*),

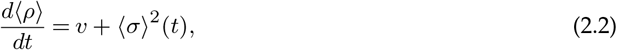

for the mean proliferation rate ⟨*ρ*⟩(*t*)

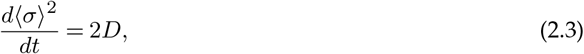

for the variance ⟨*σ*⟩^2^(*t*). These equations incorporate a constant death rate *µ >* 0, a phenotypic drift velocity *v*, which can be positive, zero or negative, and a diffusion constant *D >* 0 accounting for the fluctuations in the proliferation phenotype.

The three differential equations (2.1)-(2.3) constitute an exactly solvable model. The first one yields

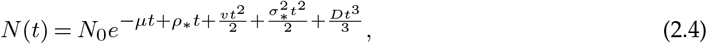

with *N*_0_, *ρ*_*∗*_ and *σ*_*∗*_ denoting the initial cell population, proliferation rate and variance, respectively. One important prediction of (2.4) is that the total cell population evolves in a *fundamentally different* fashion than a simple exponential growth *N* (*t*) = *N*_0_*e*^(*ρ∗*−*µ*)*t*^, the latter occurring if no phenotypic changes take place in the proliferation rate.

For the mean proliferation rate we find

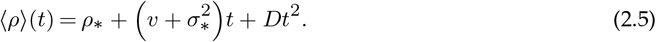

Equation (2.5) accounts for the drift in the mean proliferation seen in all of our simulations, which is quadratic with time due to the presence of the diffusion constant *D*. Notice that even in the absence of the drift velocity (*v* = 0), ⟨*ρ*⟩(*t*) still increases with time; the dominant contribution being due to the stochastic fluctuations embodied in the phenotype diffusion coefficient and, to a lesser extent, in the initial variability *σ*_*∗*_.

The time dependence of the standard deviation is

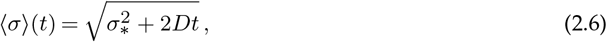

which provides another explicit and simple expression for the broadening in the phenotype landscape observed in our numerical simulations. This form for the standard deviation is characteristic of other standard diffusive processes in many biological systems [38].

Figures 2**b** (inset) and 2**c** also compare the numerical solutions of the mean proliferation rate, the standard deviation and the cell numbers obtained from our reaction-advection-diffusion model [see Eq. (4.3) in Methods] and formulas (2.4)-(2.6) giving additional confirmation of the internal consistency of our findings. Hence, in the scenario where full phenotype inheritance occurs, three distinctive features arise: a broadening in the phenotype landscape, a drift in the mean proliferation and a total cell population growing faster than a classic exponential law.

### A human non-small cell lung carcinoma line (NCI-H460) displays growth rate increase with time in-vitro

It is widely known that phenotypic plasticity is constitutively active in living organisms, including cancer [39]. However, finding an appropriate biological substrate to observe the phenotype variability is not immediate due to the coexistence of many simultaneous biological effects in most experimental models.

To test if our theoretical concepts could be acting even in simple biological scenarios we employed *in vitro* assays of human non-small cell lung carcinoma line (NCI-H460) under various culture conditions (see Methods). Real time datasets of 24 replicas with four different initial cell numbers; specifically 500, 1000, 1500 and 2000, were monitored by means of a xCELLigence Real Time Cell analyser that allowed us to estimate and track the total cell number after seeding. We then analyzed the time windows during which the mean proliferation rate exhibited an increase before cell death was detected [see Fig. 3] and competition effects could play a role in the dyamics. In all examined cases, the cell culture conditions were far from confluence.

**Figure 3.**
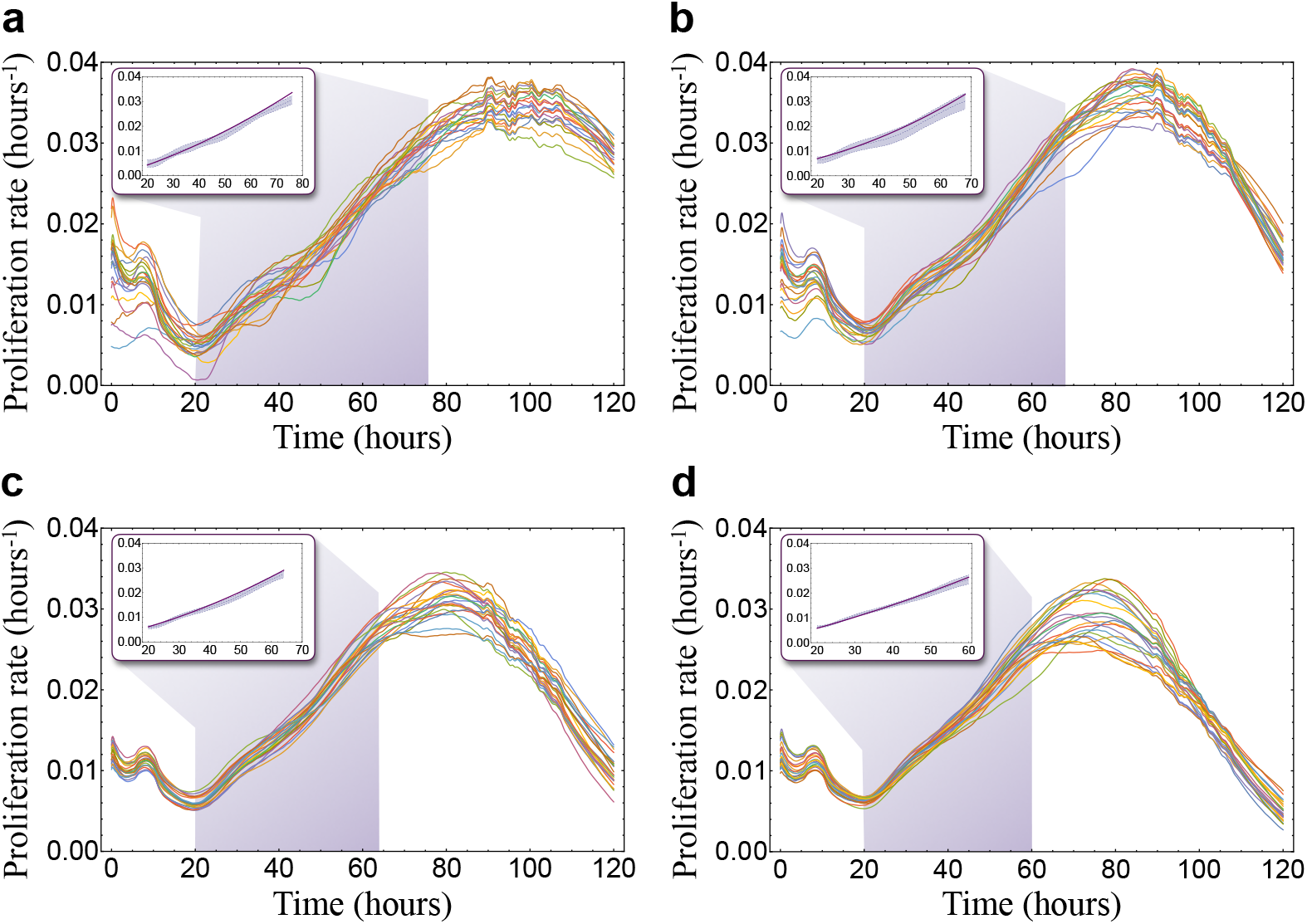
*In vitro* determination of the mean proliferation rate in a human non-small cell lung carcinoma line (NCI-H460). Proliferation rate dynamics for initial cell densities of **a** *N*_0_ = 500, **b** *N*_0_ = 1000, **c** *N*_0_ = 1500, and **d** *N*_0_ = 2000. Cell numbers were obtained using a xCELLigence Real Time Cell analyser. The number of replicas per cell density was 24. Insets show the time windows were an increase in proliferation rate was found before a substantial cell death was observed. The bands in the insets represent the experimental averaged curves, the band width corresponding to the mean standard deviation, whereas the solid curve is the fit to Eq (2.5). The obtained estimations for the phenotype diffusion coefficient were, in each case, **a** *D* = 2.61 ± 1.92 × 10^−6^ hours^−3^, **b** *D* = 4.40 ± 2.19 × 10^−6^ hours^−3^, **c** *D* = 3.37 ± 1.70 × 10^−6^ hours^−3^, and **d** *D* = 2.11 ± 2.10 × 10^−6^ hours^−3^. In all fitted cases the confidence interval for *D* was set at the level of 95%.

From the measured proliferation curve for each initial cell density and replica we fitted the data to Eq. (2.5). Figure 3 depicts the fits. A sustained increase of the proliferation rate was observed systematically in our experimental setup. Our experimental results support that the mean proliferation rate increases via a quadratic time dependence given by formula (2.5) until an observable cell death sets in, producing a net decrease in the mean proliferation. Moreover, from these results we could quantify the magnitude of the phenotypic transitions through the estimate of the phenotype diffusion coefficient, which is a key parameter in our mathematical models. The obtained estimates for the phenotype diffusion coefficient were for each initial number of cells seeded: *D* = 2.61 ± 1.92 × 10^−6^ hours^−3^ (*N*_0_ = 500), *D* = 4.40 ± 2.19 × 10^−6^ hours^−3^ (*N*_0_ = 1000), *D* = 3.37 ± 1.70 × 10^−6^ hours^−3^ (*N*_0_ = 1500), and *D* = 2.11 ± 2.10 × 10^−6^ hours^−3^ (*N*_0_ = 2000). These results suggest that phenotypic transitions sustaining the proliferation rate continuous increase could be spontaneous and take place in short time-scales even in the absence of any external selection pressures.

### Partial loss of phenotype inheritance reduces but does not eliminate average proliferation rate increase with time in-silico

We then looked at the possibility of phenotypic modifications being only transient. That is, the scenario of a *partial loss of inheritance* in the phenotypic traits by assuming that all phenotypes have a nonzero transition rate, *Γ*_*i→∗*_, of reversion to the basal phenotype with proliferation rate *ρ*_*∗*_. As in the inheritable scenario, a shift in the mean proliferation rate ⟨*ρ*⟩(*t*) towards higher values with time was observed [see Fig. 4**e**]. The magnitude of this shift was smaller as the value of *Γ*_*i→∗*_, measuring its relative relevance with respect to other processes, increased. Another visible difference with the full phenotype inheritance scenario is that now the distribution displayed a bimodal profile for a certain range of *Γ*_*i→∗*_ values, evidencing the coexistence of a peak corresponding to the clonal population distributed around the initial proliferation rate *ρ*_*∗*_ and a second broader peak comprising the more *evolved* phenotypes [see Fig. 4**a**-**d**]. The standard deviation of the phenotypic distribution [see Fig. 4**f**], ⟨*σ*⟩(*t*), increased with time for all values of *Γ*_*i→∗*_, albeit progressively less as this reverse transition rate became larger. A growth in the total cell number faster than a classic exponential still occurred [Fig. 4**g**] the magnitude of which was modulated by the partial phenotypic inheritance condition embodied in *Γ*_*i→∗*_. The larger the value of this parameter, the closer the total cell number approached a classic exponential growth.

**Figure 4.**
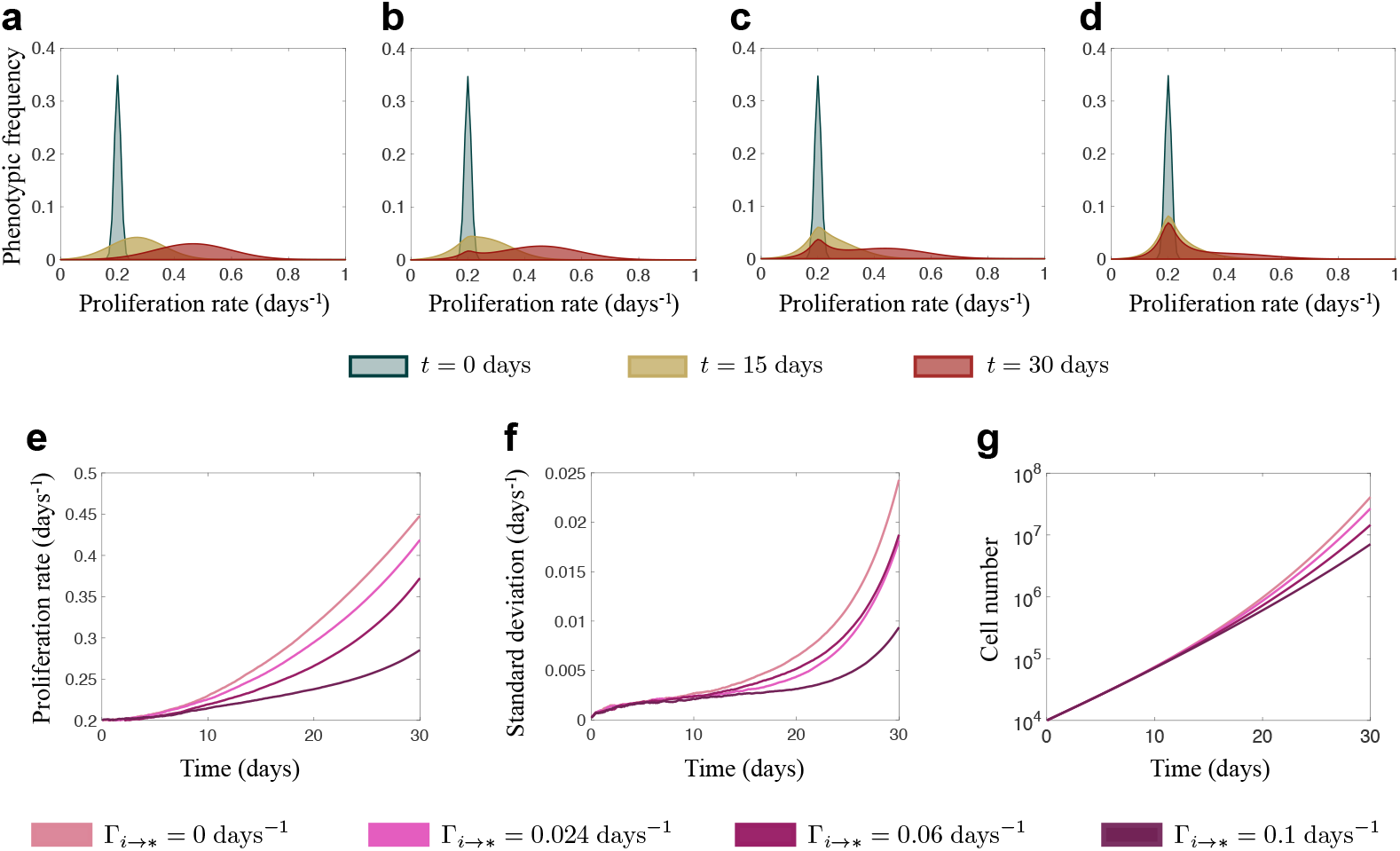
*In silico* evolutionary dynamics in scenarios of partial loss of phenotype inheritance. **a**-**d** Phenotypic frequency at times *t* = 0, 15, and 30 days for different values of the transition rate *Γ*_*i→∗*_: **a** *Γ*_*i→∗*_ = 0 days^−1^, **b** *Γ*_*i→∗*_ = 0.024 days^−1^, **c** *Γ*_*i→∗*_ = 0.06 days^−1^, **d** *Γ*_*i→∗*_ = 0.12 days^−1^. **e** Time-dependence of the mean proliferation corresponding to the cases **a**-**d. f** Time-dependence of the standard deviation corresponding to the cases **a**-**d. g** Dynamics of the total cell population corresponding to the cases **a**-**d**.

### Growth rate dynamics is not influenced by growth factors

It is reasonable to question if the shift of the average growth rate towards a higher average proliferation rate observed *in vitro* could be the result of other biological phenomena rather than a manifestation of the evolutionary mechanism proposed here. One possibility could be that proliferation could be boosted by an increasing accumulation of growth factors in the cell culture. To rule out this possibility we performed different theoretical analyses and experiments. Firstly, we implemented growth factors secretion in our discrete mathematical model. In our setting all cells secreted growth factors with the same rate (independent of their proliferation rate). The available amount of growth factors in the medium continuously enhanced proliferation in all phenotypes, slightly increasing their growth rate.

To parameterize our model, we must fix both the amount of growth factors being secreted per cell and per unit of time and to which extent growth factors influence the proliferative potential of the cell. First, average growth factor production was set to 0.1 ng cell^−1^ day^−1^ from estimates of hepatocyte growth factor (HGF) [40] and Insulin-like growth factor (IGF-1) [41] serum concentration. To estimate the quantitative influence of growth factors on growth rate we used experimental data in Ref. [42], which elucidated the relationship between epithelial growth factor (EGF) concentration and the % of centrosome separation. An estimated concentration of 200 ng was necessary to double the % of centrosome separation. In our simulations cells kept secreting growth factors with the above specified rate which accumulated in the cell culture medium afterwards. No changes in the phenotypic distribution except for a tiny shift towards a higher growth rate were observed in our simulations as shown in Supplementary Figure 1.

To study further the potential role of growth factors on the observed phenomena we put forward two additional experiments. Growth factors are typically secreted to cell surroundings, so we first conducted an experiment in cell-conditioned culture media. To do so we let cells to grow for 24h or 48h. We studied four different initial cell densities (500, 1000, 1500 and 2000 cells) in our experimental setting to assess if cell density could influence the amount of available growth factors. Cell culture medium previously conditioned by tumor cells has been formerly reported to have mitogenic activity [43]. In our experiments, cell growth did not appear to be affected by culture medium cell-conditioning [see Fig. 5a; Supplementary Figure 2 for the 24 h condition; Supplementary Figure 4 for the 48 h condition]. Phenotype difussion coefficient in the cell-conditioned medium experiments was estimated by fitting growth rate data [see Fig. 5b, see supplementary figure 3 for the 24h condition, supplementary figure 5 for the 48h condition] to equation 2.5. In general, the results revealed no statistically significant differences (*P* >0.05) [see supplementary Table 1] between the phenotype diffusion coefficients obtained for the different initial cell numbers and medium conditioning times [see Fig. 5c, Table 1].

**Figure 5.**
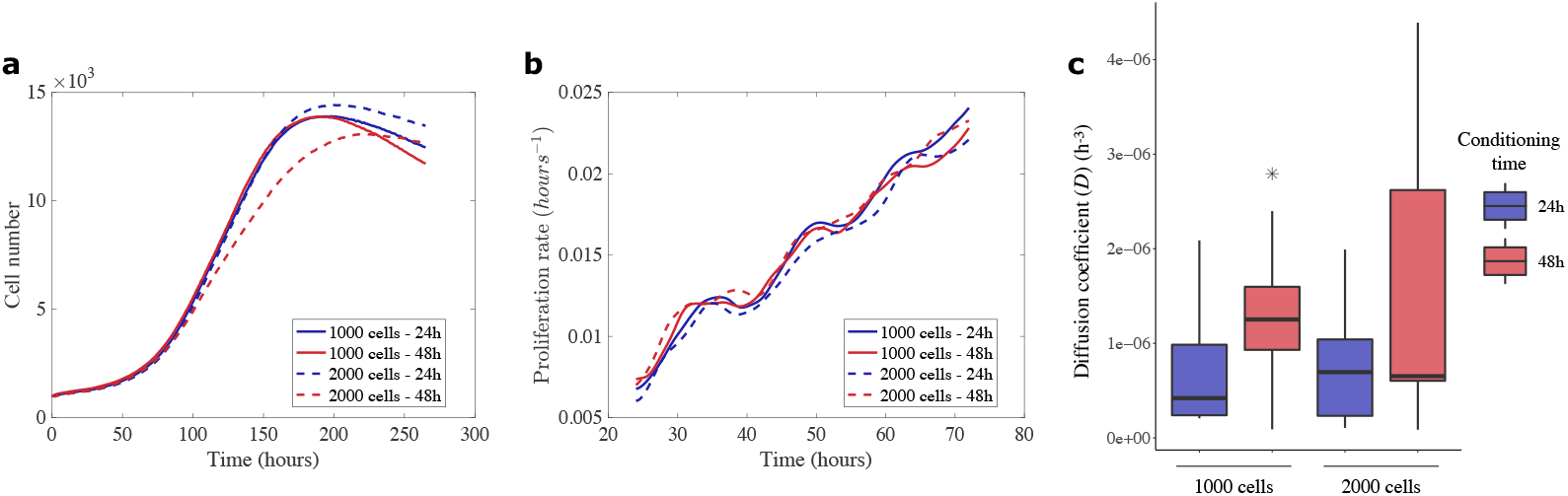
*In vitro* determination of non-small-cell lung carcinoma cells (NCI-H460) growth dynamics in the presence of previously cell-conditioned culture medium. **a** Real time assessment of non-small-cell lung carcinoma cells growth under exposure to cell-conditioned medium. Two cell densities (*N*_0_=1000 and *N*_0_ = 2000 cells) were used to produce cell-conditioned medium. Cell-conditioning was carried out over 24h (*n*_1000, 2000_ = 9) and 48h (*n*_1000_ = 9 and *n*_2000_ = 9). Average growth curves are shown for each case. **b** Growth rates for the cell number curves obtained for each experimental condition. The time derivative for *N* (*t*) was computed using a finite difference formula and then smoothed out to reduce the noise in original curves. A time window from 24 to 72 hours was analysed to prevent cell growth being affected by confluence effects. Average growth rate curves are shown. **c** Estimation of the phenotype diffusion coefficient (*D*) after fitting the growth rate curves to the mean proliferation rate explicit equation 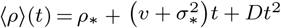. *D* values for the different groups were compared using a pairwise wilcoxon rank sum test with Bonferroni correction. No statistically significant differences were found between the different cell-conditioning setups. Outliers are represented as *.

**Table 1.** Calculation of the phenotype diffusion coefficient *D* for cell growth in cell-conditioned culture medium. Cell conditioning took place over 24 and 48 h. Four different cell densities were considered: 500, 1000, 1500 and 2000 cells. Growth rate experimental data were fitted to the explicit equation for the mean proliferation rate 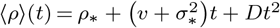 using a quadratic regression

| Incubation time<br>(hours) | Initial cell density<br>(number of cells) | D<br>(hours<sup>-3</sup>) |
| --- | --- | --- |
| 24 | 500 | $(2.26 \pm 1.18) \times 10^{-6}$ |
| | 1000 | $(1.19 \pm 7.93) \times 10^{-6}$ |
| | 1500 | $(1.57 \pm 0.76) \times 10^{-6}$ |
| | 2000 | $(1.05 \pm 5.21) \times 10^{-6}$ |
| 48 | 500 | $(2.18 \pm 1.08) \times 10^{-6}$ |
| | 1000 | $(1.30 \pm 0.76) \times 10^{-6}$ |
| | 1500 | $(1.25 \pm 0.52) \times 10^{-6}$ |
| | 2000 | $(1.04 \pm 5.21) \times 10^{-6}$ |

The second experiment monitored cell growth in the presence of a variable level of fetal bovine serum (FBS) [see Fig. 6a]. As expected, the extremely low FBS concentration of 2% considerably inhibited cell growth. However, 10% FBS and 20% FBS conditions lead to very similar results. It is worth highlighting that, despite the growth curves were influenced by the level of FBS [see supplementary figure 6], the growth rate were very similar independently of the FBS concentration [see Fig. 6b; see supplementary Figure 7]. The phenotype diffusion coefficient was found to be independent of the level of FBS [see Fig. 6c, Table 2], as revealed by the statistical analysis [see supplementary Table 2].

**Figure 6.**
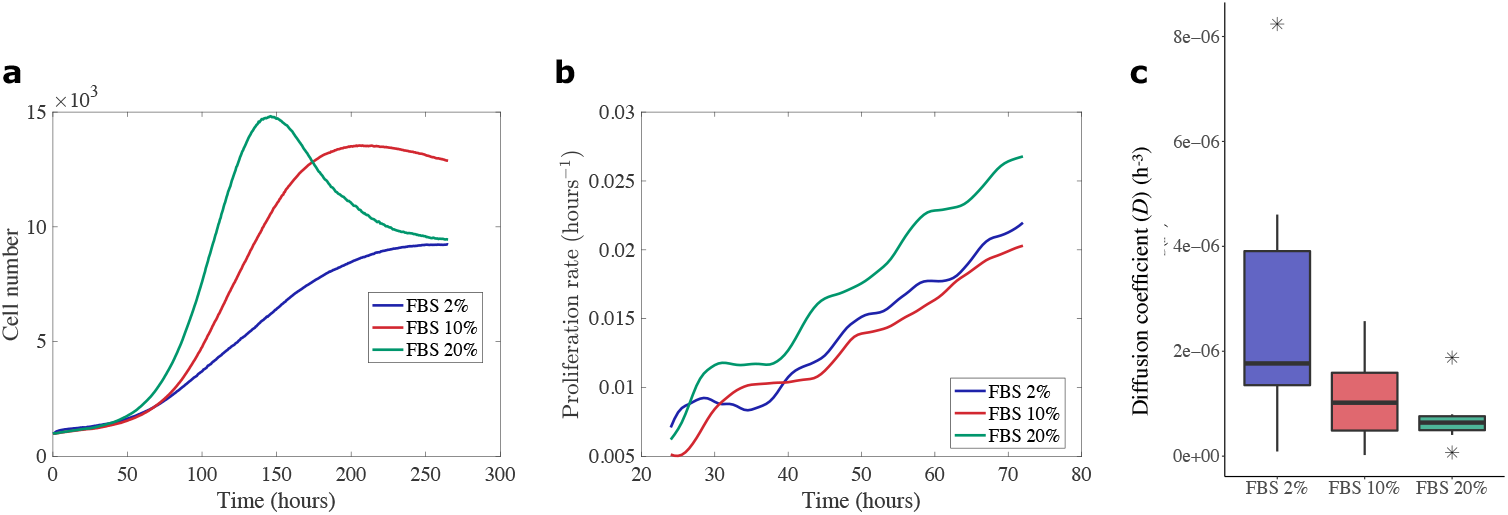
*In vitro* determination of the effect of exogenous nutrients supply or deprivation through variable fetal bovine serum (FBS) concentration on non-small-cell lung carcinoma cells (NCI-H460) cell cultures. **a** Real time assessment of non-small-cell lung carcinoma cells for initial cell density *N*_0_=1000 cells under 2% (n=6), 10% (n=8) and 20% (n=8) FBS concentration. Average curves are shown. **b** Calculation of growth rate from the cell number curves for each experimental condition. The time derivative for *N* (*t*) was computed using a finite difference formula and then smoothed out to reduce the noise in original curves. A time window from 24 to 72 hours was used to analyze the data in order to minimize effects of cell confluence. Average growth rate curves are shown. **c** Estimation of the phenotype diffusion coefficient (*D*) after fitting the growth rate curves to the mean proliferation rate explicit equation 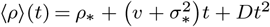. We compared *D* values using a pairwise wilcoxon rank sum test with Bonferroni correction. Outliers are represented as *. No statistically significant differences were found for the different FBS concentrations.

**Table 2.** Estimation of the phenotype diffusion coefficient *D* in NCI-H460 cell culture under variable fetal bovine serum (FBS) concentrations. Growth rate experimental data were fitted to the explicit equation for the mean proliferation rate 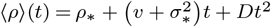 using a non-linear regression approach.

| FBS concentration (%) | Phenotype diffusion coefficient (hours<sup>-3</sup>) |
| --- | --- |
| 2 | $(3.09 \pm 2.85) \times 10^{-6}$ |
| 10 | $(1.14 \pm 8.24) \times 10^{-6}$ |
| 20 | $(1.17 \pm 1.27) \times 10^{-6}$ |

Thus, the phenotype diffusion coefficient was robust across all experimental conditions thus ruling out a potential influence of growth factors on the evolutionary dynamical phenomena reported here.

## 3. Discussion

In this study we put forward a mathematical model of an ideal cellular clonal population with stochastic changes of the cellular proliferation rates and showed that intrinsic cellular variability could lead to a sustained increase of the population average proliferation rate, even in the absence of any evolutionary pressures. We observed the same dynamics through *in vitro* real time assessment of a human non-small cell lung carcinoma line NCI-H460, what allowed us to estimate the key dynamical parameters of the mathematical model.

Tumour evolution is a complex process and nowadays it is extensively known that beyond genetic alterations, genomic aberrations, such as discordant inheritance or DNA macroalterations, and non-genetic determinants of evolution also play an important role [44]. Together, all these processes are shaping phenotypic plasticity, suggesting that deterministic dynamics could fail to explain the sequence of events observed in tumour evolution. Clonal cell lines exhibit also a higher frequency of random monoallelic expression that could increase phenotypic plasticity and spread the probability of success in a changing environment without altering the population identity [45].

For simplicity, and since we were mainly interested in characterizing how and why tumour growth accelerates in time, we focused only on proliferation. It is already known that tumour cells have a higher growth rate than proliferative non-tumoral tissues. In our study, the conceptual initial setting consisted of a genetically homogeneous clonal population, with all cells having a growth rate concentrated around an initial value. These cells were assumed to be able to slightly increase or decrease their growth rate in time with equal probability and thus explore a landscape of proliferation states while keeping a hypothetical common genotype. Interestingly, this symmetric randomness did not result in fluctuations around an average proliferation level. Instead the sole action of stochastic phenotypic transitions lead to a continuous increase of the average growth rate with time and thus to a growth of the total clonal tumour cell population fundamentally faster than the classical exponential one. This phenomenon is in line with recent findings of explosive tumour growth in different cancers [25], but it may be related to other natural contexts where dramatic increases in other population species take place, such as cyanobacteria and algae blooms occurring in eutrophic waters [46]. All these findings reveal the need for further explorations within the interplay between oncology and ecology via evolutionary theory.

Phenotypic plasticity together with noisy gene and protein levels expression has been demonstrated by means of classical and next-generation sequencing techniques such as single cell RNA-sequencing or tissue-specific differentially methylated regions (tDMRs), giving an increasing importance to the distribution of gene expression levels or epigenetics marks beyond the classical genocentric point of view [47]. These stochastic phenomena have also been reported at many other levels in nature, including differing levels of resistance to antibiotics in genetically identical bacteria or even the stochastic mechanisms underlying the development of trichromatic vision of human individual cone cells [48]. Noisy fluctuations affecting cell traits could give tumour cells the chance to sample and explore an extended landscape of physiological states, as has been reported in prokaryotes [15].

Several authors have recently analysed mathematically evolutionary cancer dynamics in phenotype-structured populations [49,50]. In those interesting works environmental selection pressures were the key issues driving phenotypic variability. Other mathematical works from a statistical mechanics perspective suggest that the number of phenotypically available states shapes tumour growth [51]. However, to the best of our knowledge, the effect of intrinsic phenotypic variability as an underlying force having a *steering* effect on the natural evolution of tumours has not been addressed in detail. While genomic instability and driver gene mutations play an essential role in the evolutionary dynamics of human cancers, the sustained increase in proliferation observed in [25], which our mathematical models also predict, has some resemblance with classic Darwinian selection ideas that are tightly linked to the selection of the fittest genotype (in our case phenotype). It is relevant to emphasize that the evolutionary dynamics observed in our models was not of Lamarckian type since we did not consider phenotypic switches to be environmentally-driven in our *in silico* models [6,52].

One of the novel aspects of this work was the spontaneous increase in average growth rate with time. A reduction in cell cycle duration over time has previously been proposed theoretically under therapy-induced cell death [36]. In that work it was assumed that a tumour population initially heterogeneous consisted of cells with different, albeit intrinsic and fixed, proliferation rates. Phenotypes were inherited and growth rate increase was the result of a Lamarckian selection process. This is different from our approach in which phenotypic diversity emerges spontaneously from a clonal population and undergoes a Darwinian evolution driven by stochastic fluctuations.

The observed increase in growth rate with time in our *in vitro* experiments could be naively attributed to the positive feedback loops affecting mitosis resulting from growth factors activity. These molecules are known to influence cell proliferation and a wide range of other cell activities including cell matrix remodelling, cell differentiation and inflammation [43]. Indeed, the studied periods of time were long enough to display the since growth factors dynamics work generally in a time scale of hours [53]. Although growth curves were affected by growth factor levels, mostly in the fetal bovine serum conditioning experiment, the phenotype diffusion coefficient remained robust across all studied experimental conditions.

Phenotype diffusion coefficient was the key mathematical parameter allowing to understand growth rate dynamics, especially for long time intervals. As it can be inferred from our explicit mathematical formulas, this parameter was involved both in the broadening of our phenotype density function *n*(*ρ, t*) and the drift of the mean proliferation rate. Our experimental results suggest that the increase of average growth rate in time in this non-small-cell lung carcinoma cell line was not linked to the activity of growth factors and other secreted molecules.

This diffusion in phenotype landscape reminds us of spreading dynamics of invasive species, in which the rate of new site colonization is not constant over time as has been proven in a variety of biological kingdoms, from viruses to vertebrates [54]. Adaptive plasticity could be considered as a trait itself and consequently be subject to evolutionary processes. However, we could expect that it does not play such an important role in a ‘constant’, non-tumour, environment. Cancer cells are exposed to varying environmental conditions across tumour development and this may lead to a short-term evolutionary response where mutational genetic variation would not be the main driving force. Phenotypic diversity is a convenient strategy for the success of population expansions in a broad range of contexts. Although it is challenging to test this kind of hypothesis at the laboratory due to the required long time-scale (as a consequence of long individual lifespan), some attempts in unicellular organisms have been reported in the literature. Those evidences reflect that at short time scales, phenotypic variations are key as a strategy to succeed in fluctuating environments as shown in *Chlamydomonas* [55] and *Lactobacillus sp*. [56], also allowing for specialization in the long-term, as shown by *Escherichia coli* culture over 2000 generations under an alternating temperature regimen [57]. The behaviour of subclonal populations interacting within the constraints of the tumour microenvironment could resemble the dynamics of the interaction of functional groups of species with variation in resource exploitation ability and environmental requirements, and the relevance of this ecological-type interactions has been evidenced even in the context of cancer [58]. Although phenotypic diversity implies an additional productivity cost for the functional group, a higher phenotypic variance seems to increase the long run performance in nature [24].

Another interesting point concerns the role of inheritance of phenotypic modifications. It is already known that phenotypic modifications in somatic cells can be passed on from one generation to another by mitosis as stated previously in the fully inheritable scenario [59,60] and they do not necessarily reverse after the inducing agent ceases [61]. In fact, increasing evidence suggest that adaptation can be graded. Short-term stresses would evoke tiny modifications in gene expression through signalling-mediated regulation of gene expression. On the other hand, a sustained stress situation could lead to a more radical switch in cell state, through epigenetic regulation or positive feedback loops, and hence drive to a permanent phenotypic modification [62–64]. We explored those different inheritance patterns in our work through the partially inheritable scenario. Our results indicated that the shift towards a more aggressive average profile in the tumour phenotypic distribution was qualitatively robust across both scenarios, even when modifying the reverse transition rate. The relationship between this shift in the proliferation phenotypes and spatial heterogeneity remains to be explored. Microenvironmental spatial heterogeneity due, for example, to the gradients of nutrients and metabolic waste generated by tumour cells [65] might also affect that reverse transition rate. The availability of physical space and new niches for dispersal and colonization might also accelerate the shift in the tumour average proliferation rate [66–68]. Indeed, the theoretical location of evolution at the tumour boundary has been previously reported [69] and the highest proliferation activity seems to be located also at the tumour edge, specially in poor prognosis cases, as it was recently underlined in two cohorts of breast and lung cancer patients [70].

The broadening in the phenotype landscape predicted by our models may play a key role not only in the heterogeneity of the clonal tumour cell population, but also in the emergence of resistance mechanisms under the administration of cytotoxic therapies [6]. Although these may successfully target the most proliferative cells, eventually those in the lower spectrum of the phenotype landscape would be able to repopulate the fastest proliferation cell states. Phenotypic diversity is a convenient strategy for the success of population expansions in a broad range of contexts. A better understanding of the fundamental biological processes underlying phenotypical plasticity as a source of intratumoral heterogeneity might be useful for tumour containment or implementing adaptive therapies [17,71], and, ultimately, for better design of therapeutic strategies.

In conclusion, in the context of mathematical models displaying phenotype plasticity, we have observed three distinctive features: a broadening in the phenotype landscape, a drift in the mean proliferation and a total cell population growing faster than classic exponential laws. Our mathematical models were conceptually simple and can be extended along many directions, including not only spatial effects (e.g. saturation), other traits besides proliferation, as well as more cell subpopulations (e.g. immune cells). The main predictions seem to be robust enough for experimental validation.

It is remarkable that these effects emerge spontaneously in the absence of selection pressures and are independent of initial cell density and phenotypic switching probability. When phenotypic traits were allowed to be lost partially, resembling the dilution of epigenetic marks as cell division progresses, the effects were still preserved. This evolution towards a more aggressive phenotype would undoubtedly be accentuated by the presence of selection pressures in the tumour microenvironment. Furthermore, we cannot ignore that this stochastic variation is probably affecting almost any cell trait and consequently tumour cell interactions with other tumour cells and also with the surrounding tissue. However, our results indicate that the existence of this stochastic non-genetic variability in the proliferation rate would be acting as permanent background force and it seems robust enough to spontaneously drive cancers to more aggressive phenotypes.

## 4. Methods

### Discrete stochastic model

We used a discrete stochastic model describing the growth dynamics of a clonal population of tumour cells having different proliferation rates, i.e. one in which not all cells divide at the same pace. To simplify the analysis, we considered a large but finite number *M* of allowed proliferation rates *ρ*_*i*_ in the interval [*ρ*_min_, *ρ*_max_]. Each cell belongs to a proliferative state *i* (with *i* = 1, 2, …, *M*) defined by its rate *ρ*_*i*_ = (*i* − 1)*Δρ* + *ρ*_min_, where *Δρ* = (*ρ*_max_ − *ρ*_min_)*/*(*M* − 1). Let *N*_*i*_(*t*) denote the number of tumour cells having phenotype *i*, thus corresponding to a rate *ρ*_*i*_, at time *t*. The total number of cells at time *t* is 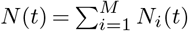. At time *t* = 0 the population, with *N*_0_ being the initial cell number, is distributed in the phenotypic landscape around a characteristic proliferation rate *ρ*_*∗*_ having a standard deviation *σ*_*∗*_. To simulate the population dynamics at later times, for every interval [*t, t* + *Δt*] in steps *Δt*, we test whether each cell has undergone a phenotypic switch, with transition rate *Γ*_*i→j*_, from proliferation state *i* to an adjacent state *j* = *i* ± 1 characterized by a proliferation rate *ρ*_*j*_. No phenotypic jumps are allowed from *i* = 1 to *j* = 0 and from *i* = *M* to *j* = *M* + 1. All these switches thus give rise to a net decrease in the number *N*_*i*_(*t*) of cells having the same *ρ*_*i*_ at time *t*. Similarly, phenotypic jumps with transition rates *Γ*_*j→i*_ from adjacent proliferation states *j* = *i* ± 1 into *i* result in a net increase in the cell number *N*_*i*_(*t*). Additionally, during time interval [*t, t* + *Δt*], mitotic and apoptotic events could also take place, each either increasing or decreasing the cell number *N*_*i*_(*t*) by one unit. Combining all these stochastic processes leads to a balance equation for the number of cells that, at time *t* + *Δt*, have a proliferation rate *ρ*_*i*_

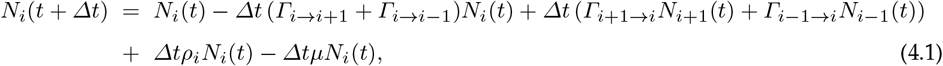

with *µ* being the death rate (taken equal for all cells). In our numerical simulations using (4.1) we assumed for simplicity that *Γ*_*i→i*+1_ = *Γ*_*i→i*−1_ = *Γ*_*i*+1*→i*_ = *Γ*_*i*−1*→i*_ ≡ *Γ*, hence giving rise to symmetric transition jumps, except at the end points *i* = 1 and *i* = *M*.

The interplay of processes described above will result in a scenario where all phenotypic changes are *inheritable*, i.e. when a cell is committed to mitosis, its progeny will be placed in the same proliferative state. This gradually yields a progressive irreversibility from the starting characteristic proliferation rate *ρ*_*∗*_ towards a different phenotypic landscape. In an alternative scenario, to explore the possibility of a *partial loss of inheritance* (and thus of partial reversibility), we also considered the effect of adding a decay probability to the starting characteristic proliferation rate *ρ*_*∗*_, equivalent to the time required to complete *m*_cd_ cell divisions. Computationally, this was implemented via a transition rate *Γ*_*i→∗*_ into a localized distribution (e.g. Gaussian) centred around *ρ*_*∗*_ for each phenotype *i*. In both scenarios (i.e., under *inheritance* or *partial loss of inheritance*), to monitor the dynamics of the population in the phenotype space, we evaluated the different population frequencies *f*_*i*_(*t*) = *N*_*i*_(*t*)*/N* (*t*), with *i* = 1, 2, …, *M*, and the mean proliferation rate 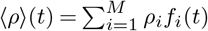.

### Continuous reaction-diffusion-advection model

To derive a partial differential equation-based model that would help to better elucidate the time dynamics of the previous discrete stochastic framework, we considered the same processes albeit we extended the phenotypic switches assuming nonnegative transition rates *Γ*_*i→j*_ from a proliferation state *i* to another state *j*, where *i, j* = 1, 2, …, *M*, and phenotypic switches with transition rates *Γ*_*j→i*_ from proliferation states *j* into *i*. Notice that 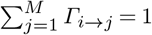 and 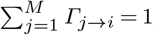, although the transition rates *Γ*_*i→j*_ and *Γ*_*j→i*_ are not necessarily equal in general. The balance equation (4.1) reads now as

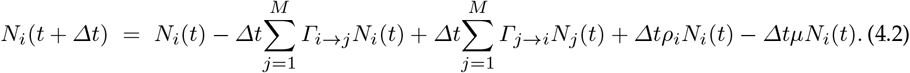

Balance equation (4.2) is quite general and encompasses the inclusion over time of new subpopulations labelled by their proliferation phenotype as their sizes become nonzero as well as the extinction of others when their cell numbers vanish. Moreover, the terms *Γ*_*i→j*_ *N*_*i*_(*t*) and *Γ*_*j→i*_*N*_*j*_ (*t*) can be understood as outward and inward cell currents for phenotype *i*, respectively.

We next perform a continuous limit approximation of (4.2). This amounts to let *Δt* → 0 and *Δρ* → 0 while the transition rates *Γ*_*i→j*_ → ∞ and *Γ*_*j→i*_ → ∞. In those limits, we assume that the two quantities 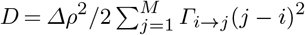 and 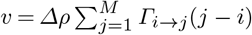 remain finite. Hence, we arrive at the following reaction-diffusion-advection equation

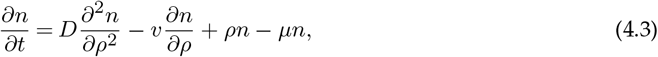

where *n* = *n*(*ρ, t*) denotes the cell density function, such that *n*(*ρ, t*) *dρ* represents the number of tumour cells that, at time *t*, have a proliferation rate between *ρ* and *ρ* + *dρ*. The first term on the right-hand side of (4.3) accounts for the fluctuations in the proliferation phenotype occurring with a diffusion constant *D* which is nonnegative. The second term describes the phenotypic drift in proliferation with a velocity *v*. Notice that this velocity may be positive or negative depending on the sign of 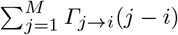 and is zero for fully symmetric or unbiased transitions. The third and fourth terms in (4.3) comprise the mitotic and apoptotic events. Additional mechanisms could be easily incorporated into (4.3), such as growth-limiting mechanisms preventing an unbounded increase in the total cell number. However, our main focus is to look at time scales for which the tumour has not yet achieved a large size.

The reaction-diffusion-advection equation (4.3) is further supplemented with initial and boundary conditions: *u*(*ρ*, 0) = *u*_0_(*ρ*) and 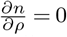, both at *ρ* = *ρ*_min_ and *ρ* = *ρ*_max_. These two zero-flux boundary conditions ensure that no cell will have a proliferation rate outside the interval [*ρ*_min_, *ρ*_max_].

### Derivation of the ordinary differential equations

The reaction-diffusion-advection equation (4.3) can be solved numerically or analytically by means of a Green’s function formalism. Here we focus on time-evolving average quantities. Specifically, the total number of cells, defined as

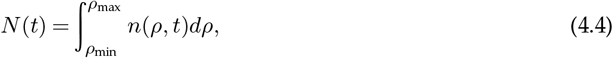

the mean proliferation rate

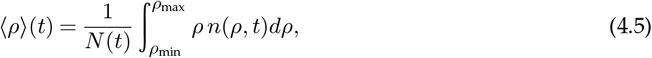

and the variance

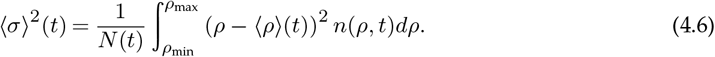

By computing the time derivative of *N* (*t*), plugging into the integrand the reaction-diffusion-advection equation (4.3), integrating by parts and imposing the boundary conditions, we get

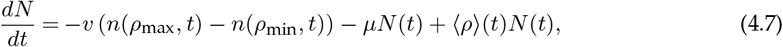

for the total cell number. As long as the cell density does not reach relevant levels at the end points of the proliferation interval, we may neglect the boundary values *n*(*ρ*_max_, *t*) and *n*(*ρ*_min_, *t*), although the term *n*(*ρ*_min_, *t*) may be nonnegligible in the case of tumour dormancy where *ρ*_min_ ≃ 0. We thus arrive at (2.1).

We may proceed in a similar fashion for the mean proliferation, and find

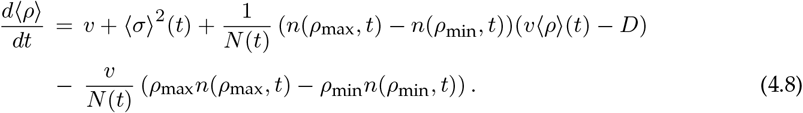

This equation reduces to (2.2) when the boundary values *n*(*ρ*_max_, *t*) and *n*(*ρ*_min_, *t*) are small. With respect to the variance, we obtain the following equation

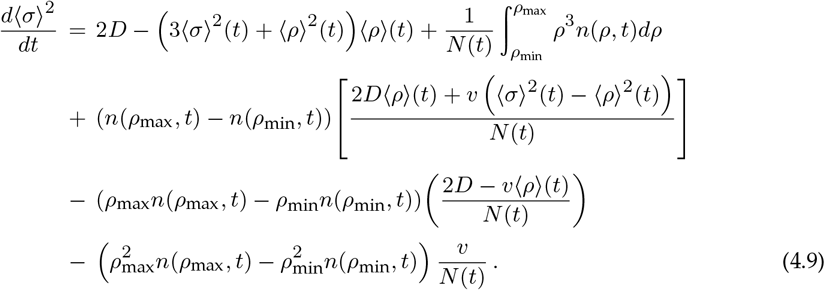

Just as in the previous derivations, we may ignore the terms involving the boundary values *n*(*ρ*_max_, *t*) and *n*(*ρ*_min_, *t*). However, the presence of the integral term on the first row of the right-hand-side of (4.9) complicates the task of obtaining a closed system of ordinary differential equations. It is nevertheless possible to obtain an approximate reduction under the stated premises by considering a Gaussian solution for the cell density of the form

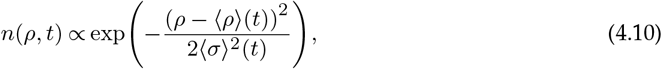

which assumes that the distribution in proliferation maintains at all times a localized profile. Using (4.10) then (4.9) simplifies to (2.3).

### Real time assessment of NCI-H460 *in vitro* growth

To experimentally study the phenotypic changes in tumour cells, we carried out *in vitro* assays under various culture conditions of human non-small cell lung carcinoma line NCI-H460 (American Type Culture Collection, Rockville, MD). The NCI-H460 cell line was maintained in a RPMI 1640 medium supplemented with 10% fetal bovine serum (FBS), 2 mM L-glutamine, and 10,000 U/ml penicillin, 10 mg/ml streptomycin, 25 g/ml amphotericin B solutions (Bioind, Beit Haemek, Israel). Cells were sub-cultured at 72 h intervals using 0.25% trypsin/EDTA and seeded into a fresh medium at 8.0 × 10^3^ cells/cm^2^ density. Continuous cell proliferation of NCI-H460 was analysed using the xCELLigence Real Time Cell analyser (ACEA Biosciences Inc., USA) which facilitates label free real-time cell analysis by measuring impedance-based signals across a series of gold electrodes. Using E-plates, 50 *µ*L of complete medium, RPMI 1640 was added to each well and the electrodes were allowed to stabilise for 30 min. The plates were then moved into the xCELLigence Real Time Cell analyser to set a base line without cells. The cells were then seeded on E-plates at four different initial number sets; specifically 500, 1000, 1500 and 2000 cells in 24 replicas per set in 0.32 cm^2^ wells in a 200 *µl* medium per well. The average maximum confluence is 32000 cells (according to the manufacturer). Cells on the electrodes were monitored by reading and recording the cell impedance every 30 min through 338 sweeps (impedance measurements started immediately). The total time duration of the cell cultures was 240 hours. The NCI-H460 cells did not exhibit contact inhibition. Their growth was only limited by available nutrients and did not undergo apoptosis during the first 80-96 hours, depending on the initial cell density.

### Estimation of cell number and mean proliferation rate from the NCI-H460 *in vitro* cell cultures

To prevent cell growth effects related to cells reaching confluence a time window from 24 to 72 hours was used to analyse the data. Cell growth data were fitted to equation (2.5) with the aim of estimating the value for the diffusion constant, *D*, associated to phenotypic transitions. The xCELLigence Real Time Cell analyser yields an arbitrary dimensionless unit called the Cell Index (CI). The CI at each time point is defined as a ratio (*R*_cells_ − *R*_back_) */*15, where *R*_cells_ is the cell-electrode impedance of the well when it contains cells, and *R*_back_ is the background impedance of the well with the media alone. The cell number *N* (*t*) and the CI are approximately related by means of a linear dependence of the form

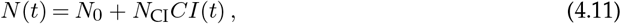

with *N*_0_ denoting the initial cell number and where in our case we calibrated the constant *N*_CI_ ≃ 1700 cells. The linearity breaks down for very low values (CI*<* 0.3) of the CI. Notice that *CI*(0) ≃ 0. We then used the CI to estimate the mean proliferation rate ⟨*ρ*⟩(*t*). Since no cell death was observed during the considered time windows (for the first 80-96 hours), we evaluated the expression

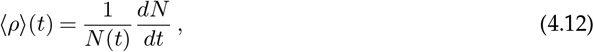

with *N* (*t*) being provided by (4.11). The time derivative of *N* (*t*) for each of the 24 replicas per initial cell number set was computed numerically using a finite difference formula and then smoothed out to reduce the presence of small noisy ripples. The numerical expressions were fitted to (2.5) under the assumption that the values for ⟨*ρ*⟩(*t*_*j*_) at different times *t*_*j*_ were independent random variables normally distributed with a common standard deviation. Since the parameters entering in formula (2.5) appear as linear combinations, we performed a linear regression analysis to estimate these parameters for each data set. Confidence intervals for the parameters were set at the level of 95%.

### Influence of growth factors on growth rate dynamics: fetal bovine serum supplementation and cell-conditioned medium

In order to evince how growth factors could be influencing growth rate dynamics, two experimental settings were proposed. On the one hand, we addressed the role of growth factor secretion through a cell-conditioned medium experiment. We employed four initial cell densities: 500, 1000, 1500 and 2000 cells, and two different incubation periods: 24h and 48h. After the incubation period, supernatants were collected for each of these sets. NCI-H460 cells were then cultured in these cell-conditioned growth media and growth was monitored using the xCELLigence Real Time Cell analyser exactly as detailed in subsection 4. A total of 9 replicates were analysed in each condition, except for the 500 cells-24h (n=8) and the 2000 cells-48h (n=7) conditions. 1000 cells were initially seeded in all cases. On the other hand, the effect of exogenous disruption of nutrients accessibility was appraised through culture medium supplementation with variable concentrations of fetal bovine serum (FBS). A starving scenario was triggered by a 2% FBS (n=6) concentration, 10% FBS (n=8) represented the standard culture conditions (control) and 20% FBS was used as a representation of nutrients overexposure (n=8). The experimental procedure for cell growth monitoring was again identical to the aforementioned (subsection 4). Statistical comparisons were performed in R (version 4.0.2) using a Pairwise Wilcoxon Rank Sum Tests with Bonferroni correction for multiple comparisons.

## Data Accessibility

Simulations were conducted in MATLAB (version R2020a). Code files for the discrete model simulations and experimental data are publicly accessible at: https://github.com/molabEvoDynamics/rep_StochasticFluctuationsDriveNonGeneticEvolution. [72]

## Authors’ Contributions

G.F.C. and V.M.P.-G. formulated the hypothesis and designed the research. J.D., A. P.-R. and M. P. conceived and performed the experiments. G.F.C., C.O.-S. and V.M.P.-G. performed the numerical simulations and statistical analyses. C.O.-S., G.F.C. and V.M.P.-G. wrote the paper. All authors gave final approval for publication.

## Acknowledgements

We thank Juan Jiménez-Sánchez and Jesús Bosque-Martínez (Universidad de Castilla-La Mancha, Spain) for the fruitful scientific discussions and Juan Antonio Delgado (University of Murcia, Spain) for discussions on ecology and evolutionary theory.

## References

1. Laland KN, Uller T, Feldman MW, Sterelny K, Müller GB, Moczek A, Jablonka E, Odling-Smee. 2015 The extended evolutionary synthesis: its structure, assumptions and predictions. Proc. R. Soc. Lond. B 282, 20151019. (doi:10.1098/rspb.2015.1019)

2. Nowell PC. 1976 The clonal evolution of tumor cell populations. Science 194, 23–28. (doi:10.1126/science.959840)

3. Cairns J. 1975 Mutation selection and the natural history of cancer. Nature 255, 197–200. (doi:10.1038/255197a0)

4. Lipinski KA, Barber LJ, Davies MN, Ashenden M, Sottoriva A, Gerlinger M. 2016 Cancer evolution and the limits of predictability in precision cancer medicine. Trends Cancer 2, 49–63. (doi:10.1016/j.trecan.2015.11.003) bibitemGreaves2015 Greaves M. 2015 Evolutionary determinants of cancer. Cancer Discov. 5, 806–821. (doi:10.1158/2159-8290.CD-15-0439)

5. Dagogo-Jack I, Shaw AT. 2017 Tumour heterogeneity and resistance to cancer therapies. Nat. Rev. Clin. Oncol. 15, 81–94. (doi:10.1038/nrclinonc.2017.166)

6. Álvarez-Arenas A, Podolski-Renic A, Belmonte-Beitia J, Pesic M, Calvo GF. 2019 Interplay of Darwinian selection, Lamarckian Induction and microvesicle transfer on drug resistance in cancer. Sci. Rep. 9, 9332. (doi:10.1038/s41598-019-45863-z)

7. Marine JC, Dawson SJ, Dawson MA. 2020 Non-genetic mechanisms of therapeutic resistance in cancer. Nat. Rev. Cancer 20, 743–756. (doi:10.1038/s41568-020-00302-4)

8. Gerashchenko TS et al. 2013 Intratumor heterogeneity: nature and biological signi?cance. Biochemistry 78(11), 1201–15. (doi: 10.1134/S0006297913110011)

9. McGranahan N, Swanton C. 2017 Clonal heterogeneity and tumor evolution: past, present, and the future. Cell 168, 613–628. (doi:10.1016/j.cell.2017.01.018)

10. Goldberg AD, Allis CD, Bernstein E. 2007 Epigenetics: a landscape takes shape. Cell 128, 635– 638. (doi:10.1016/j.cell.2007.02.006)

11. Codling EA, Plank MJ, Benhamou S. 2008 Random walk models in biology. J. R. Soc. Interface 5, 813–834 (doi:10.1098/rsif.2008.0014)

12. Huang S. 2021 Reconciling non-genetic plasticity with somatic evolution in cancer. Trends Cancer 7, 309–322. (doi:10.1016/j.trecan.2020.12.007)

13. Maheshri N, O’Shea EK. 2007 Living with noisy genes: how cells function reliably with inherent variability in gene expression. Annu Rev Biophys Biomol Struct. 36, 413–34. (doi: 10.1146/annurev.biophys.36.040306.132705)

14. Raj A, van Oudenaarden A. 2008 Nature, nurture, or chance: stochastic gene expression and its consequences. Cell. 135(2), 216–26. (doi: 10.1016/j.cell.2008.09.050)

15. Kærn M, Elston T, Blake W. 2005. Stochasticity in gene expression: from theories to phenotypes. Nat. Rev. Genet. 6, 451464. (doi: 10.1038/nrg1615)

16. Butler G, Keeton SJ, Johnson LJ, Dash PR. 2020 A phenotypic switch in the dispersal strategy of breast cancer cells selected for metastatic colonization. Proc. R. Soc. B 287, 20202523. (doi:10.1098/rspb.2020.2523)

17. Gupta PB, Pastushenko I, Skibinski A, Blanpain C, Kuperwasser C. 2019 Phenotypic plasticity: driver of cancer initiation, progression, and therapy resistance. Cell Stem Cell 24, 65–78. (doi:10.1016/j.stem.2018.11.011)

18. Byrne HM. 2010 Dissecting cancer through mathematics: from the cell to the animal model. Nature Rev. Cancer 10, 221–230.

19. Gerlee P. 2013 The model muddle: in search of tumor growth laws. Cancer Res. 73, 2407–2411.

20. Benzekry S, et al. 2014. Classical mathematical models for description and prediction of experimental tumor growth. PLoS Comput. Biol. 10, e1003800.

21. Altrock PM, Liu LL, Michor F. 2015 The mathematics of cancer: integrating quantitative models. Nature Rev. Cancer 15, 730–745.

22. Chiou SH, et al. 2008 Positive correlations of Oct-4 and Nanog in oral cancer stem-like cells and high-grade oral squamous cell carcinoma. Clin. Cancer Res. 14, 4085–4095. (doi:10.1158/1078-0432.CCR-07-4404)

23. Gupta PB, Fillmore CM, Jiang G, Shapira SD, Tao K, Kuperwasser C, Lander ES. 2011 Stochastic state transitions give rise to phenotypic equilibrium in populations of cancer cells. Cell 146, 633–644. (doi:10.1016/j.cell.2011.07.026)

24. Norberg J, Swaney DP, Dushoff J, Lin J, Casagrandi R. 2001 Phenotypic diversity and ecosystem functioning in changing environments: a theoretical framework. Proc. Natl. Acad. Sci. USA 98, 11376–11381. (doi:10.1073/pnas.171315998)

25. Pérez-García VM et al. 2020 Universal scaling laws rule explosive growth in human cancers. Nat. Phys. 16, 1232–1237. (doi:10.1038/s41567-020-0978-6)

26. Vogelstein B, Papadopoulos N, Velculescu VE, Zhou S, Diaz LA, Kinzler KW. 2013 Cancer genome landscapes. Science 340, 1546–1558. (doi:10.1126/science.1235122)

27. Huang S. 2013 Genetic and non-genetic instability in tumor progression: link between the ?tness landscape and the epigenetic landscape of cancer cells. Cancer Metastasis Rev. 32, 423– 448. (doi:10.1007/s10555-013-9435-7)

28. Batlle E, Clevers H. 2017 Cancer stem cells revisited. Nat. Med. 23, 1124–1134. (doi:10.1038/nm.4409)

29. Easwaran H, Tsai HC, Baylin SB. 2014 Cancer epigenetics: tumor heterogeneity, plasticity of stem-like states and drug resistance. Mol. Cell 54, 716–727. (doi:10.1016/j.molcel.2014.05.015)

30. Rehman SK et al. 2021 Colorectal cancer cells enter a diapause-like DTP state to survive chemotherapy. Cell 184, 226–242. (doi:10.1016/j.cell.2020.11.018)

31. Buehrlng GC, Williams RR. 1976 Growth rate of normal and abnormal human mammary epithelia in cell culture. Cancer Res. 36, 3742–3747.

32. Smith HS, Lan S, Ceriani R, Hackett AJ, Stamper MR. 1981 Clonal proliferation of cultured nonmalignant and malignant human-breast epithelia. Cancer Res. 41, 4637–4643.

33. Frick P, Paudel B, Tyson D, Quaranta V. 2015 Quantifying heterogeneity and dynamics of clonal ?tness in response to perturbation. J. Cell Physiol. 230, 1403–1412. (doi:10.1002/jcp.24888)

34. Greene JM, Levy D, Herrada SP, Gottesman MM, Lavi O. 2016 Mathematical modeling reveals that changes to local cell density dynamically modulate baseline variations in cell growth and drug response. Cancer Res. 76, 2882–2890. (doi: 10.1158/0008-5472.CAN-15-3232)

35. Powathil GG, Gordon KE, Hill LA, Chaplain MA. 2012 Modelling the effects of cell-cycle heterogeneity on the response of a solid tumour to chemotherapy: biological insights from a hybrid multiscale cellular automaton model. J. Theor. Biol. 308, 1–19. (doi: 10.1016/j.jtbi.2012.05.015)

36. Tzamali E, Tzedakis G, Sakkalis V. 2020 Modeling how heterogeneity in cell cycle length affects cancer cell growth dynamics in response to treatment. Front. Oncol. 10, 1552. (doi:10.3389/fonc.2020.01552)

37. Hanahan D, Weinberg RA. 2011 Hallmarks of cancer: the next generation. Cell 144, 646–674. (doi:10.1016/j.cell.2011.02.013)

38. Codling EA, Plank MJ, Benhamou S. 2008 Random walk models in biology. J. R. Soc. Interface 5, 813–834 (doi:10.1098/rsif.2008.0014)

39. Huang S. 2021 Reconciling non-genetic plasticity with somatic evolution in cancer. Trends Cancer 7, 309–322. (doi:10.1016/j.trecan.2020.12.007)

40. Czarkowska-Paczek B, Bartlomiejczyk I, Przybylski J. 2006 The serum levels of growth factors: PDGF, TGF-beta and VEGF are increased after strenuous physical exercise.J. Physiol. Pharmacol 57, 189–97.

41. Blum WF, Albertsson-Wikland K, Rosberg S, Ranke MB. 1993 Serum levels of insulin-like growth factor I (IGF-1) and IGF binding protein 3 re?ect spontaneous growth hormone secretion. J. Clin. Endocrinol. Metab. 76 1610–1616 (doi: 10.1210/jcem.76.6.7684744)

42. Mardin et al. 2013 EGF-induced centrosome separation promotes mitotic progression and cell survival. Dev. Cell. 25, 229–40. (doi: 10.1016/j.devcel.2013.03.012)

43. Daughaday WH, Deuel TF. 1991 Tumor secretion of growth factors. Endocrinol. Metab. Clin. 20(3), 539–63.

44. Vendramin R, Litchfield K, Swanton C. 2021 Cancer evolution: Darwin and beyond.EMBO J. 40(18): e108389. (doi: doi: 10.15252/embj.2021108389)

45. Reinius B, Sandberg R. 2015 Random monoallelic expression of autosomal genes: stochastic transcription and allele-level regulation. Nat Rev Genet. 16(11):653–64. (doi: 10.1038/nrg3888)

46. Davis TW, Berry DL, Boyer GL, Gobler CH. 2009 The effects of temperature and nutrients on the growth and dynamics of toxic and non-toxic strains of Microcystis during cyanobacteria blooms. Harmful Algae 8 715–725.(doi:10.1016/j.hal.2009.02.004)

47. Soltani M, Vargas-Garcia CA, Antunes D, Singh A. 2016 Intercellular Variability in Protein Levels from Stochastic Expression and Noisy Cell Cycle Processes. PLoS Comput. Biol. 12, e1004972. (doi:10.1371/journal.pJRSIi.1004972)

48. Geiler-Samerotte KA, Bauer CR, Li S, Ziv N, Gresham D, Siegal ML. 2013 The details in the distributions: why and how to study phenotypic variability. Curr. Opin. Biotechnol. 24, 752–759. (doi:10.1016/j.copbio.2013.03.010)

49. Ardaševa A, Anderson ARA, Gatenby RA, Byrne HM, Maini PK, Lorenzi T. 2020 A comparative study between discrete and continuum models for the evolution of competing phenotype-structured cell populations in dynamical environments, Phys. Rev. E 102, 042404. (doi:10.1103/PhysRevE.102.042404)

50. Ardaeva A, Gatenby RA, Anderson ARA, Byrne HM, Maini PK, Lorenzi T. 2020 Evolutionary dynamics of competing phenotype-structured populations in periodically ?uctuating environments. J. Math. Biol. 80, 775807.

51. West J, Newton PK. 2019 Cellular interactions constrain tumor growth. Proc. Natl. Acad. Sci. USA 116, 1918–1923. (doi:10.1073/pnas.1804150116)

52. Feinberg AP, Irizarry R A. 2010 Stochastic epigenetic variation as a driving force of development, evolutionary adaptation, and disease. Proc. Natl. Acad. Sci. USA 107, 1757–1764. (doi:10.1073/pnas.0906183107)

53. Aaronson S. 1991 Growth factors and cancer. Science. 254(5035), 11461153. (doi:10.1126/science.1659742)

54. Arim M, Abades SR, Neill PE, Lima M, Marquet PA. 2006 Spread dynamics of invasive species. Proc. Natl. Acad. Sci. USA 103, 374–378. (doi:10.1073/pnas.0504272102)

55. Reboud X, Bell G. 1997 Experimental evolution in Chlamydomonas. III. Evolution of specialist and generalist types in environments that vary in space and time. Heredity 78, 507–514. (doi:10.1038/hdy.1997.79)

56. Siezen RJ, Tzeneva VA, Castioni A, Wels M, Phan HT, Rademaker JL, Starrenburg MJ, Kleerebezem M, Molenaar D, van Hylckama Vlieg JE. 2010 Phenotypic and genomic diversity of Lactobacillus plantarum strains isolated from various environmental niches. Environ. Microbiol. 12, 758–73. (doi:10.1111/j.1462-2920.2009.02119.x)

57. Leroi AM, Lenski RE, Bennett AF. 1994 Evolutionary adaptation to temperature. III. Adaptation of Escherichia coli to a temporally varying environment. Evolution 48, 1222–1229. (doi:10.1111/j.1558-5646.1994.tb05307.x)

58. Paczkowski M et al. 2021 Reciprocal interactions between tumour cell populations enhance growth and reduce radiation sensitivity in prostate cancer. Commun Biol 4, 6. (doi:10.1038/s42003-020-01529-5)

59. Weinhold B. 2006 Epigenetics: the science of change. Environ. Health Perspect., 114, A160–A167. (doi: 10.1289/ehp.114-a160).

60. Baylin SB, Jones PA. 2016 Epigenetic Determinants of Cancer. Cold Spring Harbor perspectives in Biology, 8, a019505. (doi: 10.1101/cshperspect.a019505)

61. Turner BM. 2009 Epigenetic responses to environmental change and their evolutionary implications. Philos. Trans. R. Soc. Lond. B Biol. Sci., 364, 3403–3418. (doi: 10.1098/rstb.2009.0125)

62. Brandman O, Meyer T. 2008 Feedback loops shape cellular signals in space and time. Science. 322, 390–395. (doi: 10.1126/science.1160617)

63. Flavahan WA, Gaskell E, Bernstein BE. 2017 Epigenetic plasticity and the hallmarks of cancer. Science. 357, eaal2380. (doi: 10.1126/science.aal2380)

64. Rambow F, Marine JC, Goding CR. 2019 Melanoma plasticity and phenotypic diversity: therapeutic barriers and opportunities. Genes Dev. 33, 1295–1318. (doi: 10.1101/gad.329771.119)

65. Carmona-Fontaine C et al. 2017 Metabolic origins of spatial organization in the tumor microenvironment. Proc. Natl. Acad. Sci. USA 114, (11), 2934–2939. (doi: 10.1073/pnas.1700600114)

66. Hallastschek O, Fisher DS. 2014 Acceleration of evolutionary spread by long-range dispersal. Proc. Natl. Acad. Sci. USA 111, E4911–E4919. (doi:10.1073/pnas.1404663111)

67. Komarova NL. 2014 Spatial interactions and cooperation can change the speed of evolution of complex phenotypes. Proc. Natl. Acad. Sci. USA 111, 10789–10795. (doi:10.1073/pnas.1400828111)

68. Waclaw B et al. 2015 A spatial model predicts that dispersal and cell turnover limit intratumour heterogeneity, Nature 525, 261–264. (doi:10.1038/nature14971)

69. Deforet M, Carmona-Fontaine C, Korolev KS, Xavier JB. 2019 Evolution at the edge of expanding populations. Am. Nat. 194, 291–305. (doi:10.1086/704594)

70. Jiménez-Sánchez J et al. 2021 Evolutionary dynamics at the tumor edge reveal metabolic imaging biomarkers. Proc. Natl. Acad. Sci. USA 118, e2018110118. (doi:10.1073/pnas.2018110118)

71. Viossat Y, Noble R. 2021 A theoretical analysis of tumour containment. Nat. Ecol. Evol. 5, 826– 835. (doi:10.1038/s41559-021-01428-w)

72. Ortega-Sabater C, Fernández-Calvo G, Pérez-García VM. 2021 Stochastic ?uctuations drive non-genetic evolution of proliferation in clonal cancer cell populations. [Source code] https://github.com/molabEvoDynamics/rep_StochasticFluctuationsDriveNonGeneticEvolution

