## Supplemental figures and tables for "Stochastic fluctuations drive non-genetic evolution of proliferation in clonal cancer cell populations"

### Supplementary information

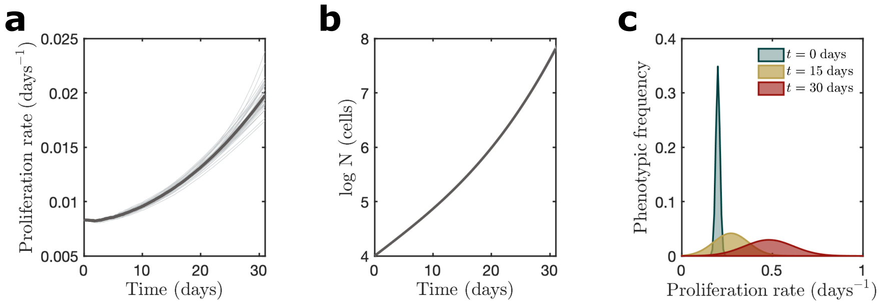


**Figure 1.** *In silico* evolutionary dynamics under the effect of the secretion of growth factors. **a** Evolution of mean proliferation rate $\langle\rho\rangle(t)$ of the distribution of $M$ phenotypes as predicted by our discrete model for $\rho \in [0,1]$ days$^{-1}$ and $t \in [0,30]$ days. Symmetric transitions jumps were considered with $\Gamma = 0.1$. Death rate ($\mu$) was equal for all phenotypes. **b** Dynamics of the total cell population for $\rho \in [0,1]$ days$^{-1}$ and $t \in [0,30]$ days. **c** Phenotypic frequency at times $t$ =0, 15, and 30 days (from left to right) of a typical run using the discrete stochastic model.

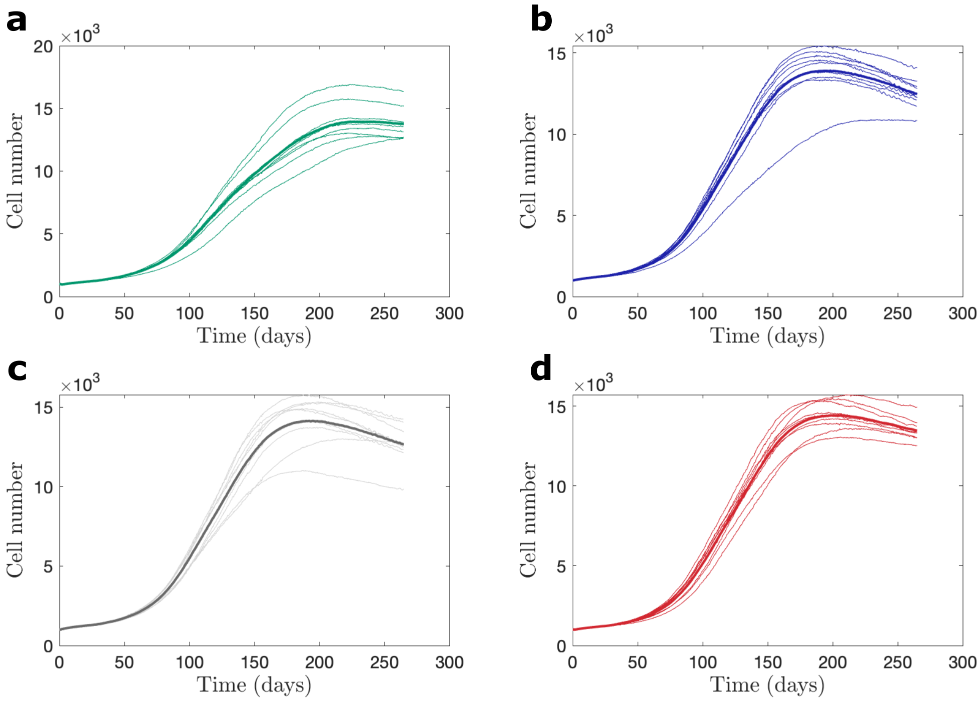


**Figure 2.** *In vitro* determination of non-small cell lung carcinoma cells (NCI-H460) in growth dynamics in the presence of previously cell-conditioned culture medium: conditioning was carried out over 24h. Each single curve accounts for an independent replicate. The average cell number is represented by the thicker line. **a** Initial seeding of 500 cells. **b** Initial seeding of 1000 cells. **c** Initial seeding of 1500 cells. **d** Initial seeding of 2000 cells.

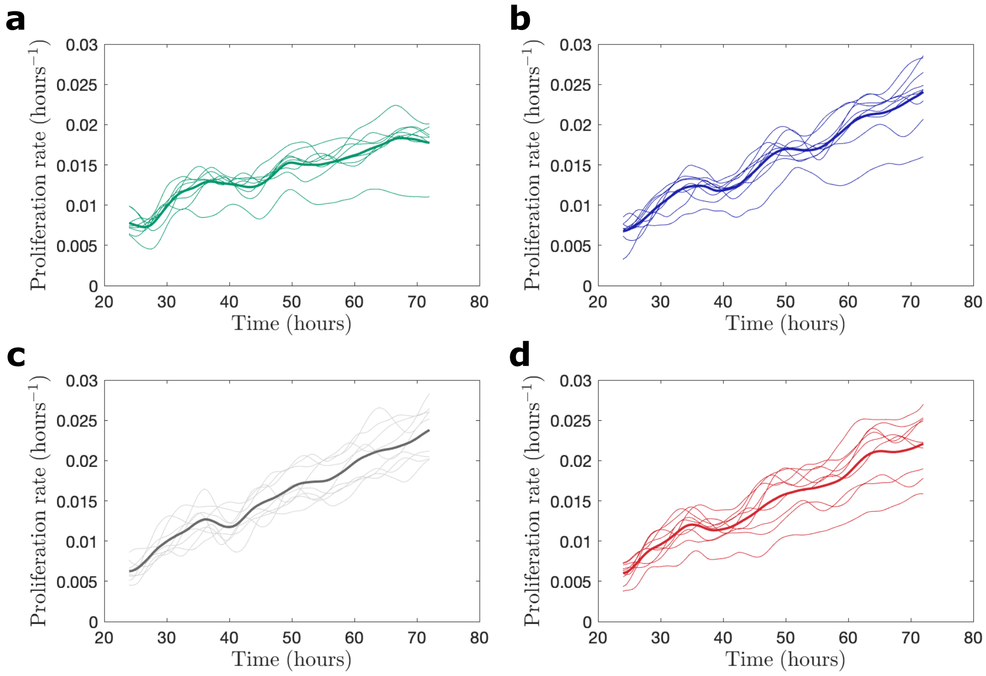


**Figure 3.** Growth rate determination of non-small cell lung carcinoma cells (NCI-H460) from collected cell number data after exposure to previous cell-conditioned culture medium: conditioning was carried out over 24h. ach single curve accounts for an independent replicate. The average cell number is represented by the thicker line. The time derivative for $N(t)$ was computed using a finite difference formula and then smoothed out to reduce the noise in original curves. **a** Initial seeding of 500 cells. **b** Initial seeding of 1000 cells. **c** Initial seeding of 1500 cells. **d** Initial seeding of 2000 cells.

| | 24h - 500 cells | 24h - 1000 cells | 24h - 1500 cells | 24h - 2000 cells | 48h - 500 cells | 48h - 1000 cells | 48h - 1500 cells |
|---|---|---|---|---|---|---|---|
| **24h - 1000 cells** | 0,434 | | | | | | |
| **24h - 1500 cells** | 0,204 | 1 | | | | | |
| **24h - 2000 cells** | 1 | 1 | 1 | | | | |
| **48h - 500 cells** | 1 | 1 | 0,152 | 1 | | | |
| **48h - 1000 cells** | 1 | 1 | 1 | 1 | 1 | | |
| **48h - 1500 cells** | 0,086 | 0,428 | 1 | 1 | 0,068 | 1 | |
| **48h - 2000 cells** | **0,034** | 1 | 1 | 1 | 1 | 1 | 1 |

**Table 1.** Statistical comparison between experimental groups under exogenous nutrients supply or deprivation through variable concentration of fetal bovine serum (FBS) in the cell culture medium. P-values resulting from the statistic analysis through a pairwise wilcoxon rank sum test with Bonferroni correction to avoid $\alpha$-error cummulation. After these correction, $P > 0.1$ are expressed like 1.

| | FBS 2% | FBS 10% |
|---|---|---|
| **FBS 10%** | 1 | |
| **FBS 20%** | 1 | 1 |

**Table 2.** Statistical comparison between experimental groups in the cell-conditioned medium experiment. All initial seeding conditions and both cell-conditioning times (24h and 48h) were tested. P-values resulting from the statistic analysis through a pairwise wilcoxon rank sum test with Bonferroni correction to avoid $\alpha$-error cummulation. After these correction, $P > 0.1$ are expressed like 1.

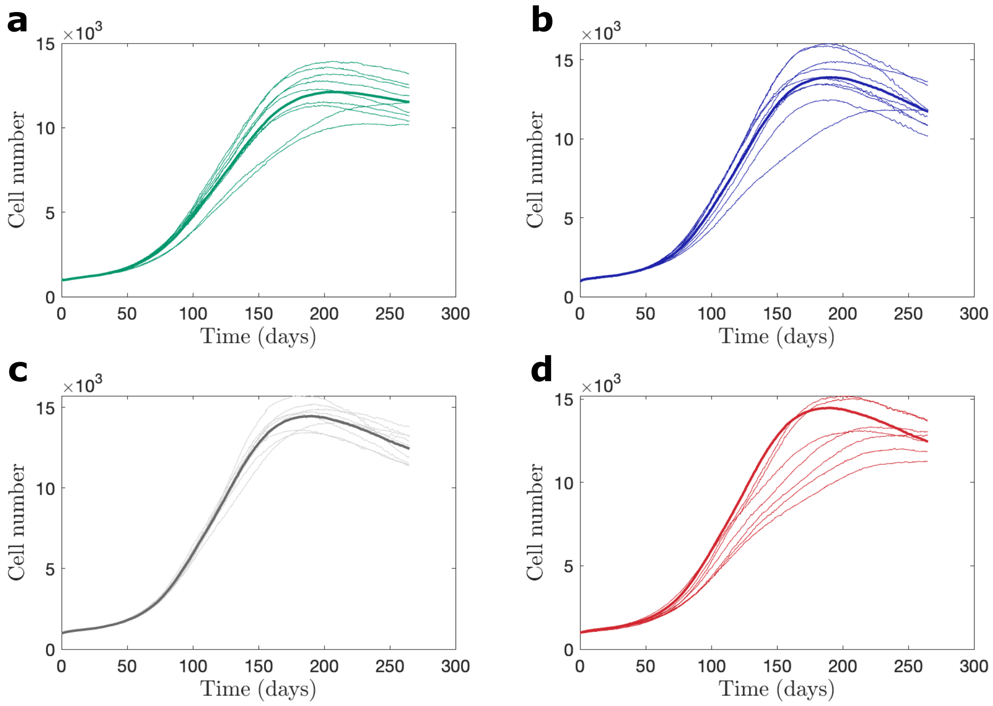


**Figure 4.** *In vitro* determination of non-small cell lung carcinoma cells (NCI-H460) in growth dynamics in the presence of previously cell-conditioned culture medium: conditioning was carried out over 48h in this case. Each single curve accounts for a replicate. The average cell number is represented by the thicker line. **a** Initial seeding of 500 cells. **b** Initial seeding of 1000 cells. **c** Initial seeding of 1500 cells. **d** Initial seeding of 2000 cells.

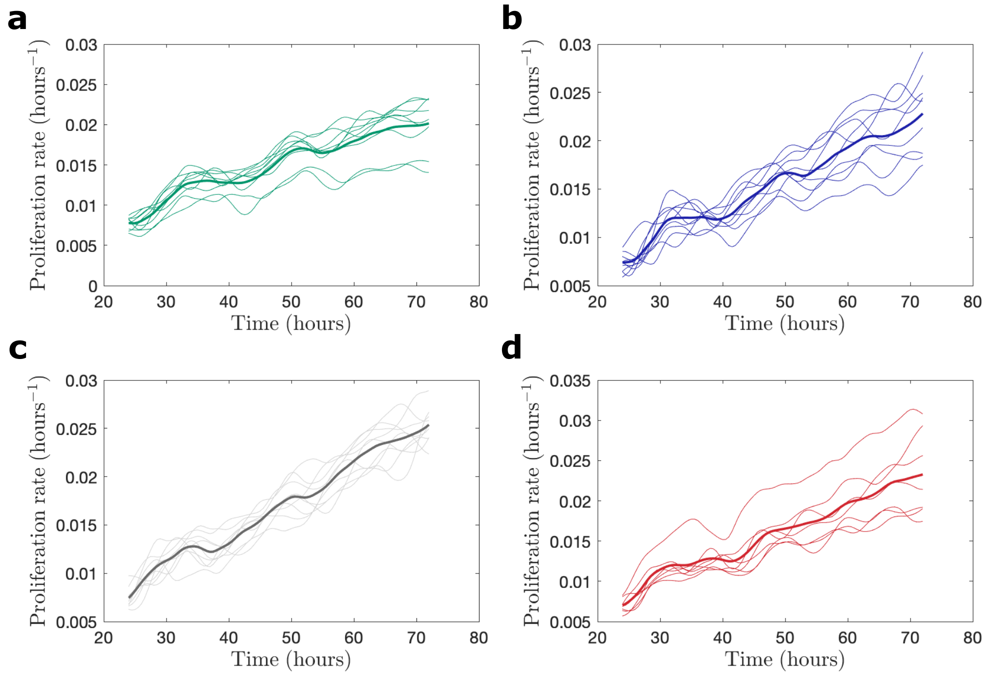


**Figure 5.** Growth rate determination of non-small cell lung carcinoma cells (NCI-H460) from collected cell number data after exposure to previous cell-conditioned culture medium: conditioning was carried out over 48h. Each single curve accounts for an independent replicate. The average cell number is represented by the thicker line. The time derivative for $N(t)$ was computed using a finite difference formula and then smoothed out to reduce the noise in original curves. **a** Initial seeding of 500 cells. **b** Initial seeding of 1000 cells. **c** Initial seeding of 1500 cells. **d** Initial seeding of 2000 cells.

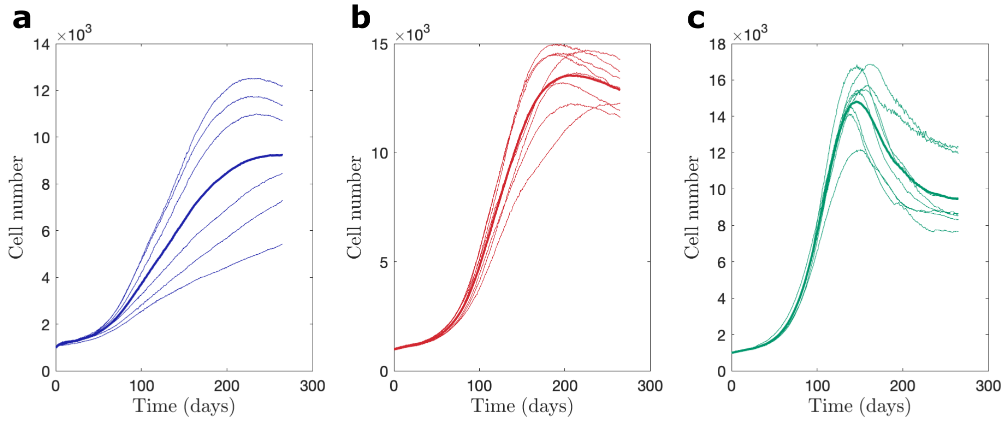


**Figure 6.** *In vitro* determination of the effect of exogenous nutrients supply or deprivation through variable fetal bovine serum (FBS) concentration on non-small cell lung carcinoma cells (NCI-H460) cell cultures. Each single curve accounts for an independent replicate. The average cell number is represented by the thicker line. Initial seeding was $N_0 = 1000$ cells. **a** Nutrients exogenous deprivation through exposure to 2% FBS. **b** Standard cell culture conditions under 10% FBS. **c** Nutrients exogenous supply through 20% FBS.

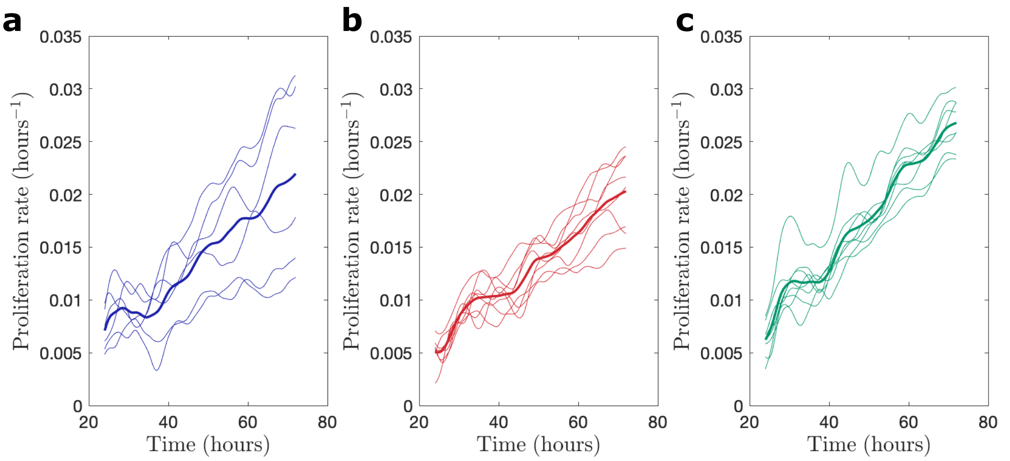


**Figure 7.** Growth rate determination of non-small cell lung carcinoma cells (NCI-H460) under the effect of exogenous nutrients supply or deprivation through variable fetal bovine serum (FBS) concentration. Each single curve accounts for an independent replicate. The average cell number is represented by the thicker line. Initial seeding was $N_0 = 1000$ cells. The time derivative for $N(t)$ was computed using a finite difference formula and then smoothed out to reduce the noise in original curves. **a** Nutrients exogenous deprivation through exposure to 2% FBS. **b** Standard cell culture conditions under 10% FBS. **c** Nutrients exogenous supply through 20% FBS.
